# Cryo-EM structure of the CDK2-cyclin A-CDC25A Complex

**DOI:** 10.1101/2023.10.17.562665

**Authors:** Rhianna J. Rowland, Svitlana Korolchuk, Marco Salamina, James R. Ault, Sam Hart, Johan P. Turkenburg, James N. Blaza, Martin E.M. Noble, Jane A. Endicott

## Abstract

The cell division cycle 25 phosphatases CDC25A, B and C regulate cell cycle transitions by dephosphorylating residues in the conserved glycine-rich motif of cyclin-dependent protein kinases (CDKs) to activate CDK activity. Here, we present the cryogenic-electron microscopy (cryo-EM) structure of CDK2-cyclin A in complex with CDC25A at 2.91 Å resolution, providing a detailed structural analysis of the overall complex architecture and key protein-protein interactions that underpin this 86 kDa complex. We further reveal an unanticipated CDC25A C-terminal helix that is critical for complex formation. Sequence conservation analysis suggests CDK1/2-cyclin A, CDK1-cyclin B and CDK2/3-cyclin E are suitable binding partners for CDC25A, whilst CDK4/6-cyclin D complexes appear unlikely substrates. A comparative structural analysis of CDK-containing complexes also confirms the functional importance of the conserved CDK1/2 GDSEID motif. This structure improves our understanding of the roles of CDC25 phosphatases in CDK regulation and may inform the development of CDC25- targeting anticancer strategies.

## Introduction

The cyclin-dependent protein kinases (CDKs) CDK1, CDK2 and CDK4/6 collectively regulate the eukaryotic cell cycle. They require association with their cognate cyclin partner for activity and are additionally controlled both by protein binding and post-translational modifications (reviewed in ^1–3^). Phosphorylation of a conserved tyrosine (equivalent to Tyr15 in human CDK1) in all eukaryotes, and additional phosphorylation of an adjacent threonine (Thr14) in metazoans, located within the glycine-rich loop (G-loop) of the ATP binding site, inhibits CDK1/2 activity. CDC25 (M-phase inducer phosphatase) was identified in *Schizosaccharomyces pombe* and shown to promote entry into mitosis by dephosphorylating CDK1 Tyr15 within the G-loop^4, 5^. In human cells, there are three CDC25 isoforms (named A, B and C^6–10^) that antagonise the activities of both Wee1 kinase that phosphorylates CDK1^11^ and CDK2 on Tyr15^12^, and Myt1 kinase that phosphorylates CDK1 on Thr14 and Tyr15^13–15^. Though early studies suggested that the CDC25 isoforms may have discrete and non-overlapping roles, the current model proposes they cooperate to regulate the major cell cycle transitions (reviewed in^16–18^). As cells transition into mitosis, CDC25B first activates CDK1-cyclin B at centrosomes in prophase^19–22^, and then all three isoforms cooperate to amplify CDK1 activity in the nucleus^23–25^. At G1/S CDC25A activity is most significant in dephosphorylating and activating both CDK2-cyclin A and -cyclin E complexes^26–28^.

CDC25 is an essential component in the response to damaged or unreplicated DNA. This role was first recognised in fission yeast, where it was demonstrated to be phosphorylated by CHK1 in response to DNA damage^29^. This pathway was subsequently found to be fundamentally conserved in mammalian cells^30, 31^. A substantial body of work has since elaborated the pathways that impact CDC25 activity and expression to regulate the cell’s response to damaged and unreplicated DNA at different stages of the cell cycle. At the molecular level, these checkpoint pathways regulate CDC25 activity through multiple mechanisms that include phosphorylation, proteolysis, binding to 14-3-3 proteins and changes in sub-cellular localisation (reviewed in^32–36^).

The majority of the post-translational modification sites that regulate CDC25 activity are located within the unstructured, extended N-terminal sequence that diverges between CDC25 isoforms and precedes their conserved C-terminal catalytic domain^37, 38^(PDB **30P3**). CDC25s are not only CDK regulators but also CDK substrates and can be recruited to cyclins A, B, D and E via an RXL motif (where X is any amino acid, CDC25A residues 11-15 RRLLF^39^ that is conserved in other CDK substrates and inhibitors (reviewed in^3^)). The association of CDC25 with 14-3-3 dimers is also mediated by residues located in the N-terminal sequence of CDC25 (reviewed in^36^). 14-3-3 binding has been proposed to directly inhibit CDC25 activity by masking the CDC25 cyclin binding motifs^40^ and indirectly by modulating its cellular location (reviewed in^41^). Phosphorylation site mutants have identified two CHK1 phosphorylation sites on CDC25A, Ser178 (N-terminal region) and Thr507 (within the C-terminal tail), that mediate 14-3-3 binding^40, 42^. The CDC25 C-terminal tail is also an important element of CDK-cyclin recognition and sequence differences between CDC25 isoforms have been shown to modulate CDC25 catalytic activity^43^.

Early studies on CDC25A and B defined them as oncogenes^44^, with changes to CDC25 expression observed in a variety of cancers (reviewed in^36, 45, 46^). Given their roles in regulating the cell cycle and response to checkpoint pathway signalling, small molecule inhibitors that target the active site have been reported (reviewed in^47^), and alternative allosteric approaches targeting CDC25-protein interactions have been considered^48^. However, there remains a lack of structural information on the binding of CDC25 proteins with their CDK-cyclin substrates, which likely hinders the development of CDC25 inhibitors and CDC25-targetting anticancer strategies. Whilst the catalytic domain structures of CDC25A^37^, CDC25B^38^ and CDC25C (PDB **3OP3**) have been solved, a structure of any CDC25 isoform in complex with a CDK-cyclin module is yet to be elucidated.

Here, we present the cryogenic electron microscopy (cryo-EM) structure of CDK2-cyclin A in complex with CDC25A at 2.91 Å global resolution. Together these proteins constitute an ∼ 86 kDa complex, making it one of the smallest asymmetric particles to have been solved by cryo-EM to a resolution better than 3 Å. This structure reveals the overall binding mode of CDC25A with CDK2-cyclin A, detailing interactions between the CDC25A catalytic domain and the N- and C-terminal lobes of CDK2. We identify a previously unobserved CDC25A C-terminal helix that binds at the CDK2-cyclin A interface and is critical for trimeric complex formation. Sequence conservation analysis against the cell-cycle CDKs and cyclins (CDK2 vs 1, 3, 4 and 6 with cyclin A vs B, E, D1 and D3) suggests CDK1/2-cyclin A, CDK1-cyclin B and CDK2/3-cyclin E are suitable binding partners for CDC25A, whereas CDK4/6-cyclin D sequences diverge at key interaction sites and appear unlikely substrates for CDC25A. These results support a model in which CDK4 and CDK6 are not regulated by G-loop phosphorylation or are dephosphorylated by other means. Comparisons with existing structures of Thr160-phosphorylated CDK2 (pT160CDK2) bound to kinase-associated phosphatase (KAP^49^) and to cyclin-dependent protein kinase subunit 1 (CKS1^50^) indicate the conserved GDSEID motif (single letter amino acid code, residues 205-210 in CDK2) is an mutual binding site for these proteins that also imparts selectivity, highlighting its importance as a CDK regulatory site.

## Results and Discussion

### Sample preparation and preliminary data collection

To prepare the CDK2-cyclin A-CDC25A complex (phosphorylated on CDK2 Tyr15 and Thr160, and hereafter called CDK2-cyclin A-CDC25A), proteins were expressed in *E. coli* cells and purified separately by affinity, ion-exchange and size exclusion chromatography (SEC). To prolong complex engagement, catalytically inactive CDC25A was prepared by mutating Cys431 to serine, and doubly phosphorylated pY15pT160CDK2 was generated by co-expression with the human Wee1 kinase catalytic domain and subsequent *in vitro* phosphorylation with *S. cerevisiae* CDK-activating kinase (CAK). The purified proteins were incubated, and the trimeric complex was isolated by analytical SEC before application to cryo-EM grids (Supplementary Figure 1).

Preliminary data collection on a 200 kV Glacios readily yielded promising 2D class averages, (Fig. 1a). However, significant preferential orientation was evident (Fourier space coverage ∼10% estimated by Cryo-EF^51^) that limited 3D refinement (Supplementary Figure 2a-c). Particle picking with Topaz^52, 53^ improved the number of unique 2D views, (Fig. 1a-b), yielding a reconstruction that encompassed the full complex. However, the map exhibited considerable directional resolution anisotropy, visualised by streaking in the EM density (Supplementary Figure 2d-f).

**Figure 1:**
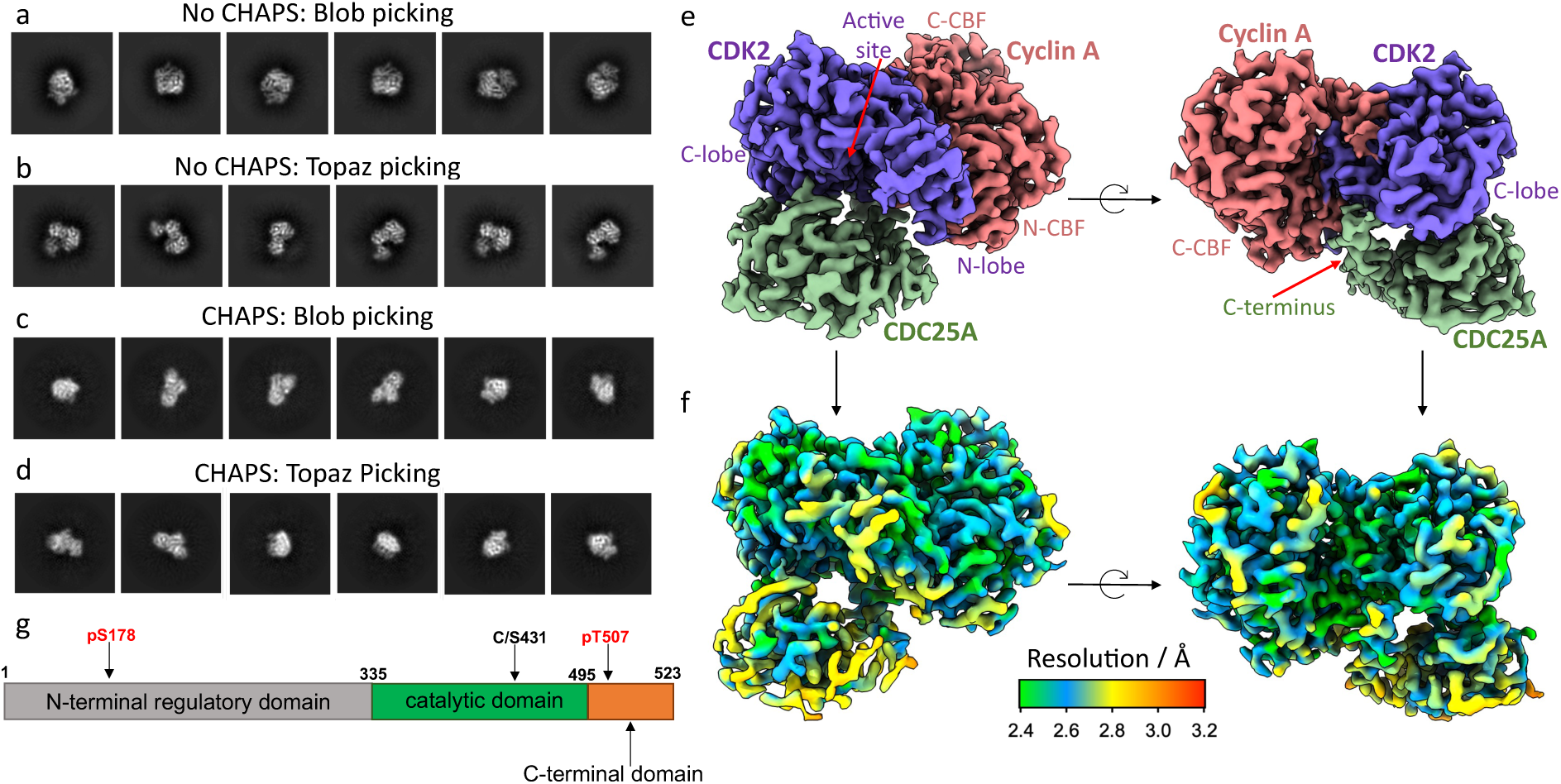
Cryo-EM structure of the CDK2-cyclin A-CDC25A complex. (a-c) Representative 2D class averages of particles of the CDK2-cyclin A-CDC25A complex picked from preliminary data using (a) blob picking, showing most 2D classes represented a front-on view of CDK2-CDC25A and (b) Topaz picking, which improved the number of unique 2D views, but preferential orientation persisted. (c) During high-resolution data collection, the use of CHAPS resulted in a wider variety of 2D class averages from (c) blob picking and (d) Topaz picking. (e-f) Cryo-EM map of the CDK2-cyclin A-CDC25A complex at 2.91 Å global resolution (EMD-**18470**). (e) CDC25A (green) catalytic domain binds across the N- and C-terminal lobes of CDK2 (purple), whilst the newly identified C-terminal helix (indicated by the red arrow) binds at the CDK2-cyclin A (salmon) interface. N- and C-terminal cyclin box folds are indicated by N- and C-CBF respectively. (f) The local resolution of the complex ranges 2.4 to 3.2 Å (FSC 0.143) (map shown in same orientations as panel e). (g) Schematic of the domain organisation of CDC25A; the catalytic domain and C-terminal tail were expressed in this study with mutation of catalytic Cys341 to serine. Phosphorylation sites that mediate the interaction with 14-3-3 proteins are labelled. For clarity, the 16 reported phosphorylation sites within the N-terminal regulatory domain (listed in Uniprot entry P30304 and annotated in Sur et al.^36^) are omitted.

To improve the particle orientation distribution, the complex was spiked with CHAPS at concentrations ranging 0.1 – 1.0 x CMC (critical micelle concentration) immediately prior to grid preparation. The grids were screened on a 200 kV Glacios and the CHAPS concentration most beneficial for improving the particle orientation was evaluated from the 2D class averages. The positive effect of CHAPS was evident at all concentrations tested; however, a wider variety of unique 2D views were identified using 0.5 and 1.0 X CMC CHAPS (Fig. 1c-d).

### High-resolution data collection and structure determination

High-resolution data collection on grids prepared with 0.5 and 1.0 X CMC CHAPS was performed on a 300 kV FEI Titan Krios equipped with a BioQuantum K3 detector and GIF energy filter (Supplementary Table 1). The full data processing workflow is described in Supplementary Figure 3, but in summary, particles of the CDK2-cyclin A-CDC25A complex were picked using blob picker, 2D classified and sorted into 3D classes. Particles from the best 3D class were chosen to train Topaz for further picking^52^. Topaz-picked particles were 2D and 3D classified, with the best 3D class refined by homogeneous and non-uniform refinement to yield the final reconstruction at 2.91 Å resolution constituting 670,852 particles. The resulting EM map (EMD-**18470**) showed clear density for the overall quaternary architecture, secondary structure, and side chain organisation of the trimeric complex (Fig. 1e and Supplementary Figure 4).

ModelAngelo^54^ was used for automated model building of the trimeric complex, completed by manual re-building of low-confidence regions in Coot^55^ according to the EM map, followed by real-space refinement in Phenix^56, 57^ to generate the full CDK2-cyclin A-CDC25A model (PDB **8QKQ**). The local resolution of the complex ranges between 2.4 Å and 3.2 Å (Fig. 1f), with the highest resolution within the CDK2-cyclin A core, and the lowest resolution within the N-terminal β-sheet domain of CDK2 (residues 9-18) and flexible loop regions of CDC25A (residues 420-424, and 492-495). Nevertheless, all expressed residues for each protein (human CDK2 residues 1-298, bovine cyclin A2 residues 169-430 Uniprot P30274 (equivalent to human cyclin A residues 171-432, Uniprot P20248), and human CDC25A residues 335-524 (Fig. 2g)) could be modelled.

**Figure 2:**
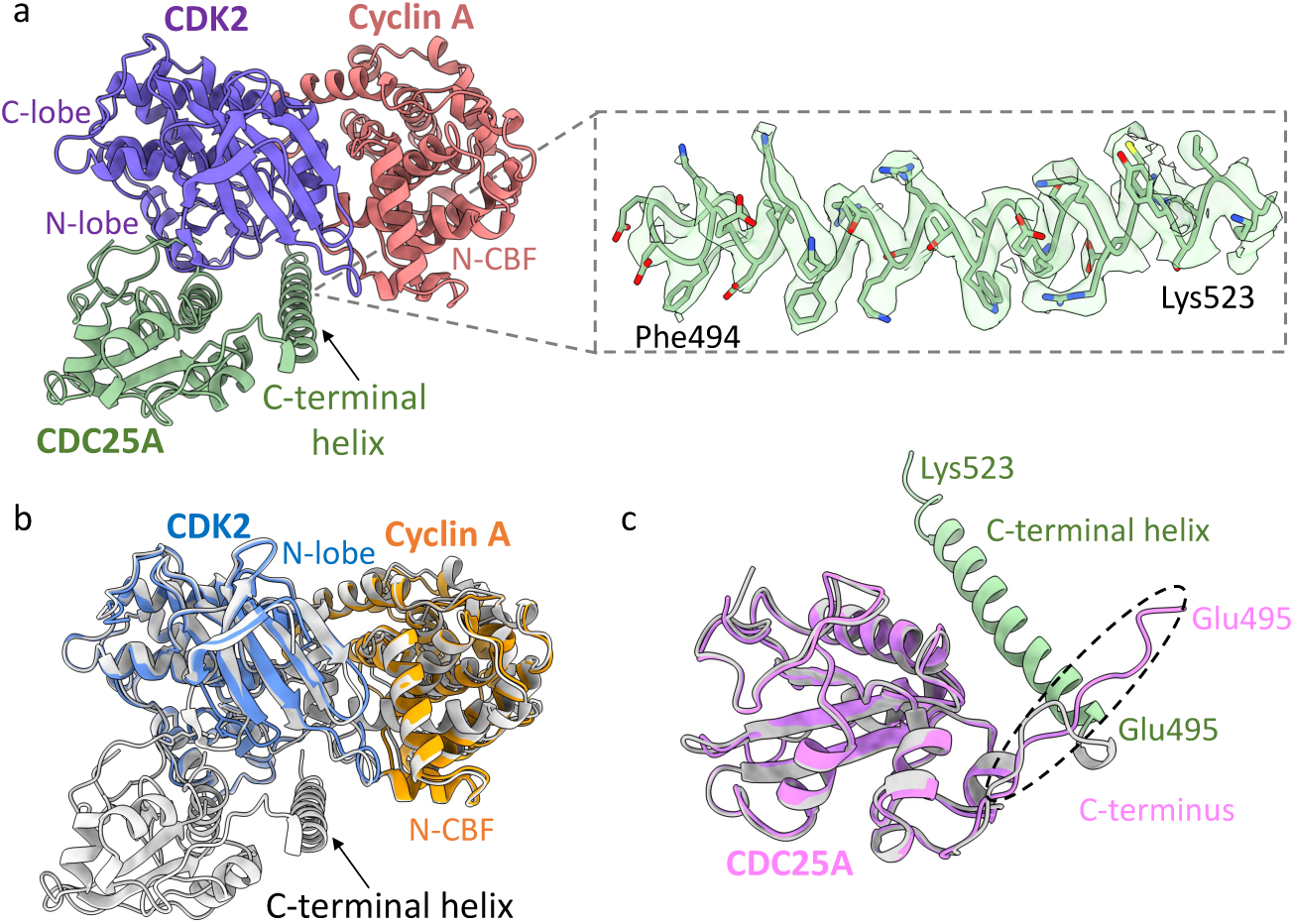
Model of the CDK2-cyclin A-CDC25A complex. (a) Structure of the CDK2-cyclin A-CDC25A complex (PDB **8QKQ)** in the same orientation and colour depiction as Fig. 1e, showing overall quaternary architecture and binding of CDC25A. Zoomed panel depicts the CDC25A C-terminal helix (EM density threshold 0.248) which binds at the CDK2-cyclin A interface. (b-c) Superposition of CDK2-cyclin A-CDC25A (grey) with existing (b) CDK2-cyclin A crystal structure (PDB **1JST**^58^, CDK2 in blue and cyclin A depicted in orange) and (c) CDC25A crystal structure (PDB **1C25**^37^, pink). The helical secondary structure of the CDC25A C-terminal domain, spanning residues Glu495-Lys523, is not observed in the CDC25A crystal structure which terminates at Glu495 (labelled in (c)).

### Overview of the CDK2-cyclin A-CDC25A complex

In the trimeric complex, CDK2 has its characteristic bi-lobal structure, comprising the β- sheet N-terminal and predominantly helical C-terminal lobes, both of which are generally well conserved with that of CDK2 in the CDK2-cyclin A complex (PDB **1JST**^58^, Fig. 2a-b). The structure of cyclin A is extremely well conserved, consisting of two helical domains (the N- and C-terminal cyclin box folds, N-CBF and C-CBF) essentially identical in chain topology that bind both lobes of CDK2 to form an extensive protein-protein interface (Fig. 2a-b).

The CDC25A catalytic domain structure is broadly consistent with the monomeric crystal form (PDB **1C25A**^37^), comprising an α/β-domain with a central parallel β-sheet enclosed by 5 α-helices (residues 335-524) (Fig. 2c). The mutated catalytic residue (Cys431Ser) initiates the conserved PTP C(X)_5_R loop between a central β-strand and α-helix, creating a shallow active site that recognises CDK2 pTyr15. The CDC25A catalytic domain bridges the bi-lobal structure of CDK2, binding on the opposite face to cyclin A, to form an extensive but discontinuous interface of ∼ 1260 Å^2^ (Fig. 2a). In contrast to the CDC25A crystal structure which visibly terminates at Glu495, the CDC25A sequence can be visualised to Lys523 (Fig. 2c). This longer C-terminal tail coverage reveals a helix, spanning residues 495-523, which binds at the CDK2-cyclin A interface relatively distant from the CDC25 core (Fig. 2a). This helix interacts with both CDK2 and cyclin A, providing a mutual binding site in the trimeric complex. Specifically, the CDC25A C-terminal helix interacts with the N-terminal αC helix (PSTAIRE, residues 45-51) and C-terminal activation segment (residues 145-172 between the DFG and APE motifs and includes phosphorylated Thr160) of CDK2, whilst providing the only point of contact for cyclin A. The interaction with the C-terminal cyclin box fold (C-CBF) of cyclin A is relatively remote, forming a small protein-protein interface of ∼ 170 Å^2^. The lack of electron density for these C-terminal residues in the existing CDC25A crystal structure, indicates this region of CDC25A may be disordered, transitioning to nascent secondary structure on binding the CDK2-cyclin A substrate.

We further analyzed the CDK2-cyclin A-CDC25A complex in solution by hydrogen-deuterium exchange mass spectrometry (HDX-MS), comparing monomeric CDC25A against the trimeric complex (Supplementary Figure 5). This analysis highlighted a region of CDC25A that is significantly protected on complex formation (Supplementary Figure 5a-c); specifically peptide 1 (VRERDRLGNEYPKLHYPEL, residues 445-463) which starts towards the middle of the helix C-terminal to the mutated catalytic Cys431Ser residue and continues into the succeeding loop. The VRERDRLGNEY and PKLHYPEL portions of peptide 1 show ∼10% and ∼15% difference in fractional uptake respectively. Although the latter portion of this peptide sequence (residues 457-464, PKLYHPEL) is partially protected in the monomeric form by the most N-terminal CDC25A residues, the enhanced protection for the full peptide sequence (VRERDRLGNEYPKLHYPEL) in the trimeric complex can be rationalised by its interaction with the CDK2 GDSEID motif (described in detail below), which binds above the C-terminal end of this central CDC25A helix. Supporting this CDK2-CDC25A interaction site, reciprocal HDX analysis of CDK2 (CDK2-cyclin A vs the trimeric complex) (Supplementary Figure 5d-f) revealed significant protection for CDK2 residues 196-220 (MVTRRALFP**<u>GDSEID</u>**QLFRIFRTLG), which include the GDSEID sequence that makes important interactions with CDC25A (detailed below).

Notably, the HDX-MS data did not reveal any significant protection for the C-terminal domain of CDC25A, which is surprising given the dominant feature of the CDC25A C-terminal helix in the cryo-EM map. Given the relatively distant nature of the interaction with the CDK2-cyclin A interface (the CDC25A C-terminal helix backbone sits ∼ 11 Å away from the backbone structure of CDK2), it is hypothesized that the C-terminal helix is not substantially solvent-protected in the bound state. Additionally, the structurally dynamic behaviour of this C-terminal region in solution may result in rapid exchange in the unbound state; therefore, it may not be possible to detect binding on the timescales measured by HDX-MS.

### CDK2-CDC25A interactions: CDC25A substrate recognition

Substrate recognition by CDC25A is driven by the CDK subunit, and this interaction is mediated through three distinct CDK2 regions; the G-loop (connecting the β1 and β2 strands of the N-terminal lobe, residues 9-18 containing pTyr15), the activation segment and the GDSEID-αG helix motif. Across the CDK2-CDC25A interface, multiple hydrogen bonds and salt bridges were identified by PISA (protein interfaces, surfaces and assemblies) analysis^59^.

Within the CDC25A active site, clear density for CDK2 pTyr15 was observed (Fig. 3a-c). Although density for the surrounding CDK2 residues was more ambiguous, the backbone structure for all residues of the G-loop could be modelled. In dimeric pT160CDK2-cyclin A (PDB **1JST**), the G-loop is pulled towards the surface of CDK2, placing Tyr15 within hydrogen bonding distance of Glu51, and underneath the triphosphate of the ATP substrate. In the dimeric pY15pT160CDK2-cyclin A structure (PDB **2CJM**^60^) the Tyr15 phosphate group is solvent exposed and is coordinated through a network of waters to Ser46 and Thr47 at the start of the αC-helix^60^. When in complex with CDC25A, this G-loop is drawn away from the surface of CDK2, positioning pTyr15 into the shallow active site of CDC25A (Fig. 3a-b). In this extended loop conformation, pTyr15 hydrogen bonds with the mutated catalytic Ser431 residue and surrounding Glu432, Ser435, Glu436 and Arg437 residues of CDC25A (Fig. 3c).

**Figure 3:**
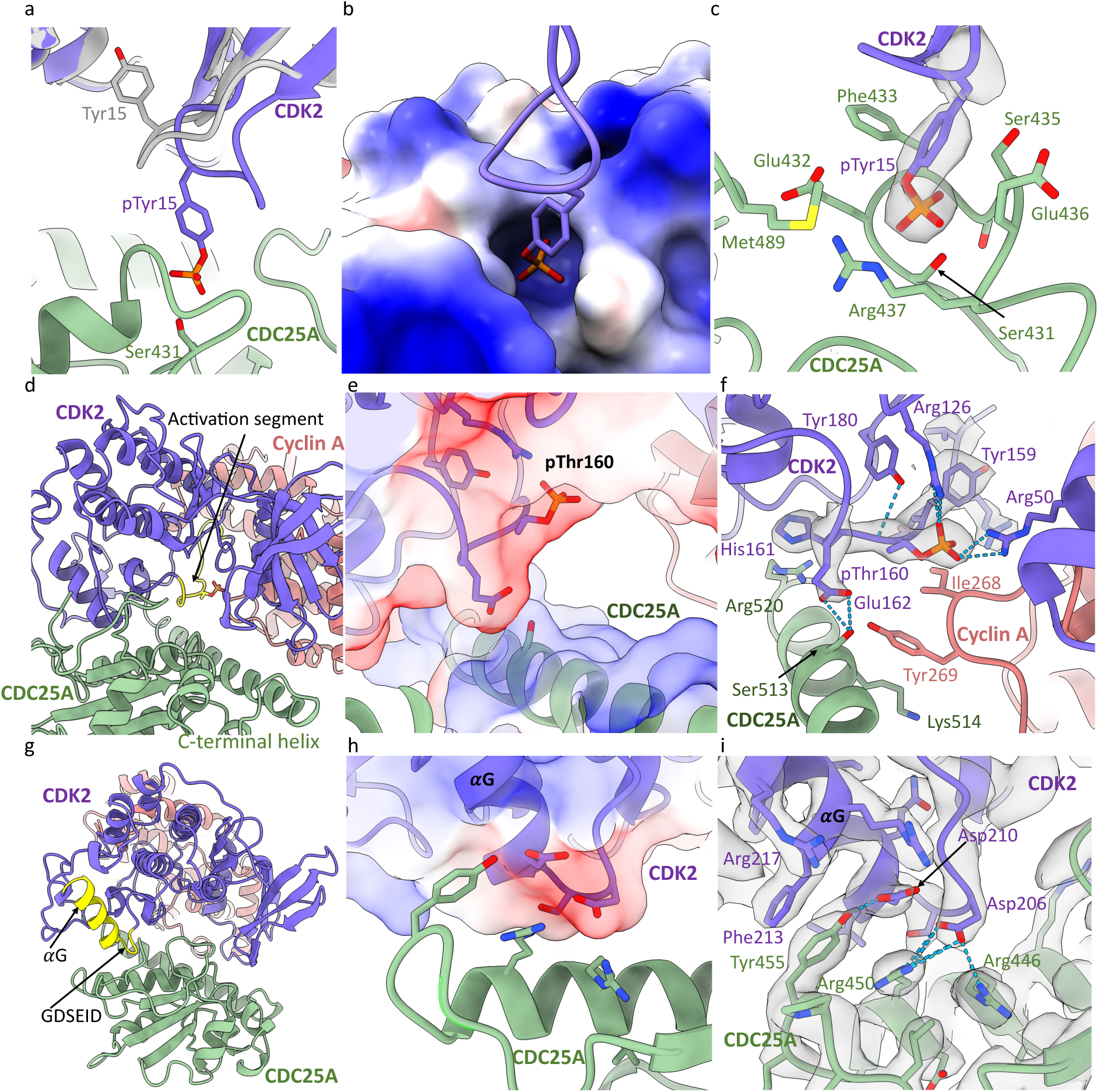
Interactions of CDC25A with CDK2. (a-c) Interactions of the CDK2 G-loop with the CDC25A active site. (a) Overlay of CDK2 (purple) with pT160CDK2-cyclin A crystal structure (PDB **1JST,** grey), highlighting a change in G-loop conformation. (b) Electrostatic surface of CDC25A active site, showing binding of CDK2 pTyr15. (c) Ribbon and density diagram (0.193 threshold) for the binding of CDK2 pTyr15 in the CDC25A active site. (d-f) Interactions of the CDK2 activation segment with the CDC25A C-terminal helix. (d) Ribbon diagram of the trimeric complex, highlighting the CDK2 activation segment (yellow). (e) Electrostatic surface of the CDK2 activation segment and CDC25A C-terminal helix. (f) Ribbon and density diagram (threshold 0.227) depicting binding of the CDK2 activation segment with the CDC25A C-terminal helix, which also binds cyclin A Tyr269 (salmon) (g-i) Interactions of the GDSEID motif with CDC25A. (g) Ribbon diagram highlighting the CDK2 GDSEID motif and αG helix (yellow). (h) Electrostatic profile of CDK2 GDSEID motif and (i) Ribbon and density diagram (threshold 0.332) for binding of the CDK2 GDSEID motif (purple) with the central helix of CDC25A (green).

In contrast the CDK2 activation segment (containing pThr160) remains relatively unperturbed by CDC25A binding, adopting a conformation almost identical to that reported in the pT160CDK2-cyclin A crystal structure (PDB **1JST**). Importantly, this conformation facilitates interaction with the CDC25A C-terminal helix (Fig. 3d-f). Unambiguous density for pThr160 shows the phosphate group resides in a small, positively charged pocket within the CDK2 activation segment and hydrogen bonds with surrounding CDK2 Arg126 and Arg150 residues. Analysis of the electrostatic surface potential suggests the activation segment provides a negatively charged surface that binds the opposingly charged surface of the CDC25A C-terminal helix (Fig. 3e). A significant interaction in this region is a hydrogen bond formed between CDK2 Glu162 and CDC25A Ser513, whilst neighbouring CDK2 His161 contacts CDC25A Arg520 (Fig. 3f). The cyclin A Tyr269 hydroxyl group (equivalent to human cyclin A Tyr271) is also within hydrogen bonding distance of CDC25A Ser513 and is placed to engage in a π stacking interaction of its aromatic ring with the aliphatic sidechain of CDC25A Lys514 within the C-terminal tail (see below). Tyr269 and Ile268 further create a notable hydrophobic patch on the cyclin A surface. Collectively, these interactions with the CDC25A C-terminal helix provide a mutual binding site for CDK2 and cyclin A within the trimeric complex.

Within the C-terminal lobe of CDK2, the αG helix and preceding GDSEID sequence protrude over the CDC25A helix that lies C-terminal to the mutated catalytic Cys431Ser residue. This generates an interface where the GDSEID motif provides a negatively charged surface which binds the positively charged CDC25A helix (Fig. 3h). At this interface, CDK2 Asp206 (the first Asp of the GDSEID sequence) hydrogen bonds to CDC25A Arg450 and forms a salt bridge with CDC25A Arg446 (Fig. 3i), whilst the backbone of Glu208 hydrogen bonds with the side chain of CDC25A Glu447. Additionally, Asp210 (the second Asp of the GDSEID motif) hydrogen bonds to nearby CDC25A Tyr455, which itself sits within hydrogen bonding distance to CDK2 Arg217 further along αG. In this region, the aromatic side chain of CDC25A Tyr455 also forms a ρε-stacking interaction with CDK2 Phe213 (Fig. 3i), which binds in a hydrophobic pocket contributed by CDK2 residues Ile209, Phe216, Phe240, Pro241 and Trp243. Combined, these residues of the αG helix and GDSEID sequence (Asp206, Glu208, Asp210, Phe213 and Arg217) create interactions that pin the CDC25A catalytic core to the C-terminal domain of CDK2.

Additional regions of CDC25A-CDK2 interaction include an extended CDC25A loop (residues 350-358) that binds in the wide groove between the CDK2 N-terminal β-sheet and the C-terminal L12-αF-L13 sequence prior to the GDSEID motif. This interaction bridges the N- and C-terminal lobes of CDK2, where CDK2 Lys129, Gln131, Thr165, Trp167, Tyr168 create a pocket that binds CDC25A Gln355. Specifically, CDK2 Gln131 hydrogen bonds with CDC25A Gln355, whilst CDC25A Lys353 hydrogen bonds to CDK2 Lys88 (Fig. 4a). Other interacting CDK2 residues are located on the strand that connects the N-terminal β-sheet to the PSTAIRE (αC) helix (Fig. 4b). This strand forms a generally negatively charged environment that binds the positively charged CDC25A C-terminal helix. Most notably, CDK2 Thr39 forms a hydrogen bond with CDC25 Arg502 and Glu42 form a salt bridge with CDC25A Arg506.

**Figure 4:**
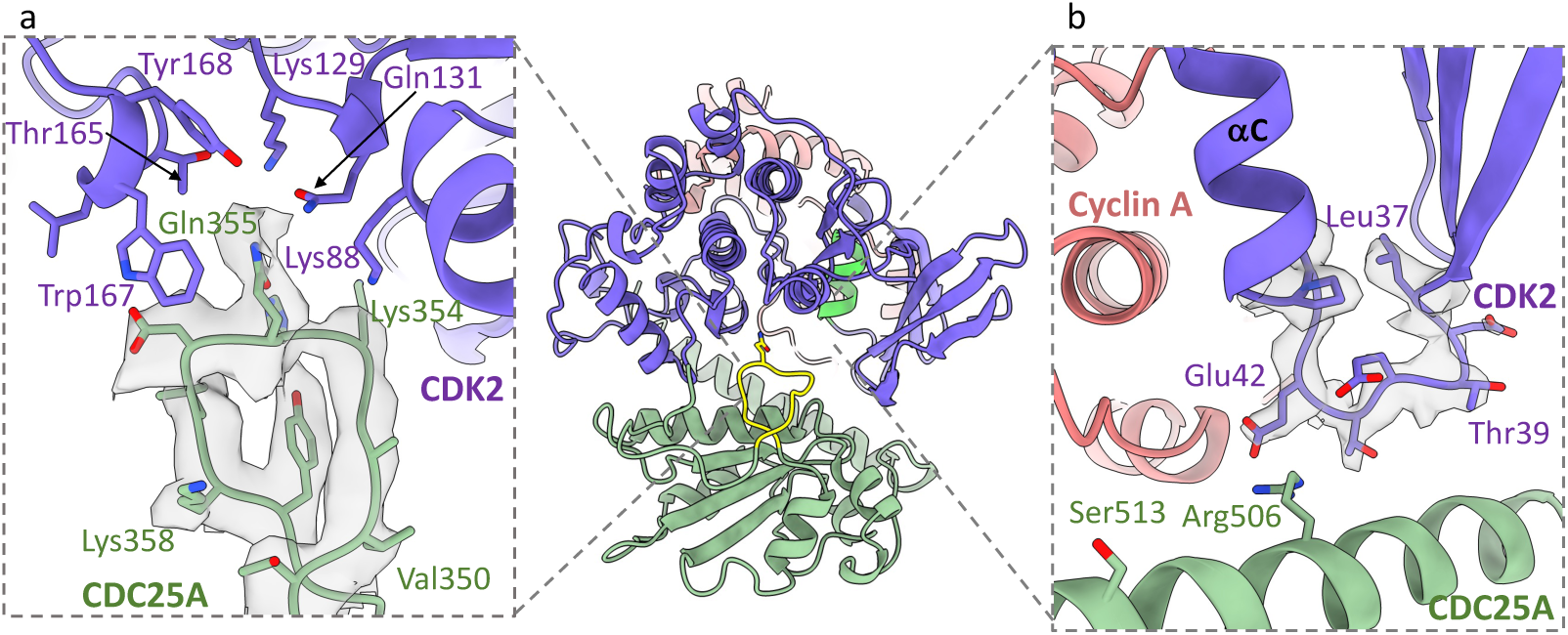
Additional CDK2-CDC25A interactions. Ribbon and density diagram (threshold 0.214) depicting (a) the binding of the CDC25A loop (residues 350-358 green) in the groove between the N- and C-terminal lobes of CDK2 (purple) and (b) binding of the CDK2 loop (residues 37-45, before the PSTAIRE (αC) helix, purple) with the C-terminal helix of CDC25A (green).

### Investigating cyclin A-CDC25A interactions

In contrast to the numerous interactions with CDK2, CDC25A makes relatively few contacts with cyclin A; only 2 hydrogen bonds and 1 salt bridge were identified between the two proteins, all of which occur across the previously unobserved CDC25A C-terminal helix (Fig. 5a-c). At the end of the central helix (α3) in the cyclin A C-CBF, Tyr269 (Tyr271 in the human cyclin A2 sequence) creates a ρε-aliphatic side chain interaction with CDC25A Lys514, whilst hydrogen bonding to CDC25A Met517 and CDC25A Ser513 which also binds the activation segment of CDK2 (Fig. 5b-c). Further N-terminal, bovine cyclin A Glu272 (equivalent to human cyclin A2 Glu274) forms a salt bridge with CDC25A Arg506 (Fig. 5c), a residue that also interacts with CDK2 through the PSTAIRE (αC) helix. Therefore, CDC25A Arg506, Ser513 and Lys514 play pivotal roles in securing the binding of the C-terminal helix to both cyclin A and CDK2. Moreover, the CDK2-cyclin A interface forms a generally negatively charged groove that binds the opposingly charged CDC25A C-terminal helix (Fig. 5b).

**Figure 5:**
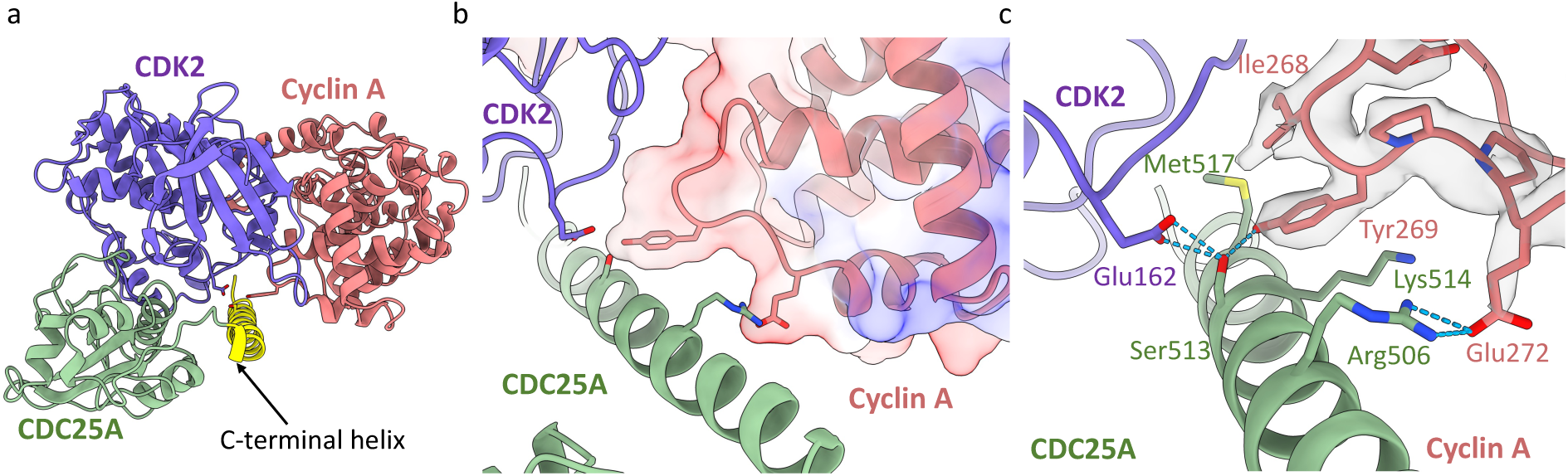
Interactions of the CDC25A C-terminal helix with cyclin A. (a) Ribbon diagram of the trimeric complex (CDK2 purple, cyclin A salmon, CDC25A green) revealing the binding of the CDC25A C-terminal helix (highlighted in yellow) at the CDK2-cyclin A interface. (b) Electrostatic surface profile of cyclin A C-CBF. (c) Ribbon and density diagram (threshold 0.278) depicting the binding of the CDC25A C-terminal helix (green) with cyclin A C-CBF. CDC25A Ser513 binds both cyclin A Tyr269 and CDK2 Glu162.

### The CDC25A C-terminal helix mediates complex formation

The CDC25A C-terminal sequence has previously been shown to mediate interaction with 14-3-3 proteins and we confirmed its requirement for binding to CDK2-cyclin A by assaying a set of CDC25A C-terminal tail mutants through homogenous time-resolved fluorescence (HTRF) (Supplementary Figure 6). As predicted from the interactions described above, terminating the CDC25A construct at Glu495, and charge reversal mutations in the CDC25A C-terminal helix (R502E/K504E/R506E and K514E/R520E) abrogated binding to CDK2-cyclin A, highlighting the importance of the C-terminal helix for trimeric complex formation.

14-3-3 proteins recognise CDC25A phosphorylated on Thr507 as one of the two phosphoamino acids required for their bidentate ligand interaction. In our cryo-EM structure, Thr507 points out into solution and, as expected, analysis by HTRF revealed mutation of Thr507 to a glutamate had little effect on the affinity of CDC25A for CDK2-cyclin A. To be accommodated within the 14-3-3 phospho-peptide recognition cleft, we hypothesise that the C-terminal region of CDC25A must be structurally dynamic, likely disordered in solution to promote recognition by the CHK1 catalytic site and subsequent 14-3-3 binding but forming helical secondary structure on binding to the CDK2-cyclin A substrate.

### The CDK2 GDSEID sequence mediates CDK2-protein interactions

Within the CDK C-terminal lobe, the CDK family has a loop linking the αF and αG helices that contains the GDSEID sequence and is conserved in CDKs 1-3. The importance of this motif to CDK1 function was first recognised through the isolation of multiple *Schizosaccharomyces pombe cdc2* cell cycle mutants (compiled in Endicott et al.^61^). This GDSEID motif is not conserved in other CDKs (Fig. 6a) and is extended by 9 residues in CDK8 (residues 240-248). The structures of CDK2 bound to cyclin-dependent kinase subunit (CKS)1 (PDB **1BUH**^50^) and kinase associated phosphatase (KAP) (PDB **1FQ1**^49^) were the first to illustrate the importance of this region to CDK1/2 regulation.

**Figure 6:**
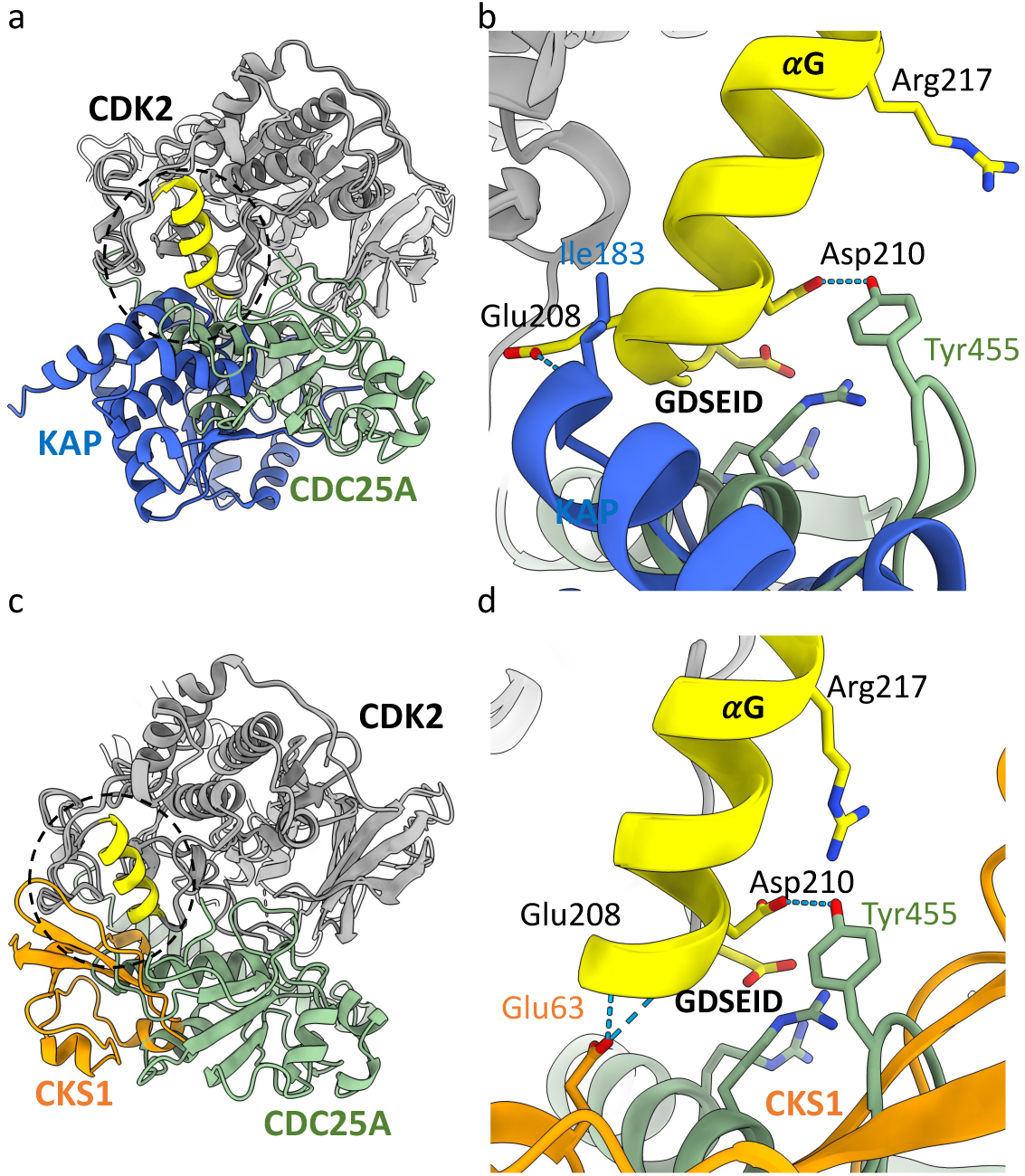
Comparison of KAP/CKS1/CDC25A with GDSEID binding region of CDK2. (a) Overlay of the trimeric complex (CDK2-Cyclin A in grey, CDC25A in green) with the CDK2-KAP crystal structure (PDB **1FQ1**, KAP in blue) and (c) CDK2-CKS1 crystal structure (PDB **1BUH,** CKS1 in orange). The CDK2 GDSEID sequence and αG helix are highlighted in yellow. (b) Ribbon diagram of the GDSEID region of CDK2 bound to CDC25A (green) and KAP (blue), which bind to opposing faces of the αG helix. (d) Ribbon diagram of the GDSEID region of CDK2 bound to CDC25A (green) and CKS1 (orange), showing an overlap in the binding region but differences in specific residue interactions.

### Comparison with the pT160CDK2-KAP complex

KAP is a dual specificity phosphatase that dephosphorylates pThr160 within the CDK2 activation segment. Superposition of the crystal structure of KAP bound to pT160CDK2 (PDB **1FQ1**) with our ternary pY15pT160CDK2-cyclin A-CDC25A complex, reveals that although the two phosphatase binding sites overlap, there are significant differences between the interfaces.

As described above, CDC25A binds across the N- and C-terminal lobes of CDK2, whereas KAP binds almost exclusively to the C-terminal lobe (Fig. 6a). In complex with KAP, the CDK2 activation segment adopts an extended conformation so that pThr160 can reach the KAP active site. However, in complex with CDC25A, the activation segment sits closer to the CDK2 surface. Therefore, CDC25A and KAP appear to require and/or induce different activation segment conformations. The two phosphatases also induce different G-loop conformations. When bound to CDC25A, this loop extends away from the CDK2 surface so that pTyr15 is placed in the CDC25A catalytic site; whereas bound to KAP it adopts a more compact conformation, where the only notable interaction occurs between KAP Lys54 and CDK2 Tyr15.

The GDSEID, αG helix and the DYK motifs in the L14 loop of CDK2 (residues 235-237) contribute affinity and specificity to the CDK2-KAP interaction^49^. The inner face of the αG helix and GDSEID sequence creates a non-polar pocket that binds residues from the C-terminal helix of KAP, where CDK2 Glu208 hydrogen bonds to KAP Ile183 (Fig. 6b). In complex with CDC25A, the outer face of the αG helix provides a negatively charged surface that binds a positively charged CDC25A core helix through hydrogen bonding interactions with CDK2 Asp206, Asp210 and Arg217 (Fig. 3i and 6b). Therefore, although the GDSEID-αG helix motif provides a mutual binding site, CDC25A and KAP bind to opposite faces of the αG helix and interact with different CDK2 residues.

### Comparison with the pThr160CDK2-CKS1 complex

CKS1 is an essential CDK accessory protein that in all eukaryotes promotes the multisite phosphorylation of CDK1 substrates by providing a phospho-amino acid binding site *in trans*^62^. In higher eukaryotes it is also an essential adaptor protein that regulates p27KIP1 abundance by facilitating its interaction with the SCF-SKP2 E3 ligase for ubiquitin-mediated proteasomal degradation^63, 64^. The CDK GDSEID motif is also required for CKS1 binding.

Whilst the recognition sites for CDC25A and CKS1 (PDB **1BUH**) partially overlap at the GDSEID region, the exact nature of the interactions is different, with the binding of CKS1 being more comparable to that of KAP. Specifically, one loop of CKS1 (residues 58–64) protrudes into a groove formed by CDK2 residues Ile209, Phe213, Phe240, Pro241, and Trp243, whilst the anti-parallel β-sheet core of CKS1 packs against the GDSEID-αG helix (Fig. 6c). This forms an extensive hydrophobic interface that spans the length of the αG helix, in which CKS1 Glu63 and Ile59 hydrogen bond with CDK2 Glu208 and Lys237 in the L14 loop that follows αG (Fig. 6d). In contrast, CDC25A interacts predominantly with the GDSEID sequence and the start of αG, forming a smaller, more hydrophilic interface strengthened by hydrogen bonds to CDK2 Asp206, Asp210 and Arg217. Therefore, the binding of CKS1 extends further across the GDSEID-αG helix and the interactions appear more hydrophobic.

On comparison of the KAP, CKS1 and CDC25A complexes, it is evident that the CDK2 GDSEID motif is an important mutual binding region for these proteins which confers selectivity by engaging different residues and hydrophobicity surface profiles. That it is a hotspot for protein binding provides the mechanistic explanation for the identification of multiple yeast cell cycle mutants within the sequence. The *S. pombe* mutants DL2^65^ (Glu215Lys, equivalent to CDK2 Glu208) and *cdc2-E8*^66^ (Asp213Asn, equivalent to CDK2 Asp206) only block at the G2/M transition with reduced kinase activity and are here shown to be at residues required for the interaction with CDC25. Additionally, the N-terminal CDK2 binding regions, notably the G-loop and PSTAIRE helix, are vital to the binding of CDC25A, whilst these regions appear less important for the binding of KAP and are irrelevant for CKS1 binding, providing further affinity and selectivity for CDC25A.

### Sequence alignments to identify CDC25 substrate preferences

This cryo-EM complex provides a template from which to assess the sequence differences between members of the CDK, cyclin and CDC25 families (Table 1, Supplementary Figure 7) to identify potential partner preferences. In addition to CDKs 1, 2 and 3, CDK4 has been reported to be phosphorylated on the conserved tyrosine within the G-loop^67^.

**Table 1:**
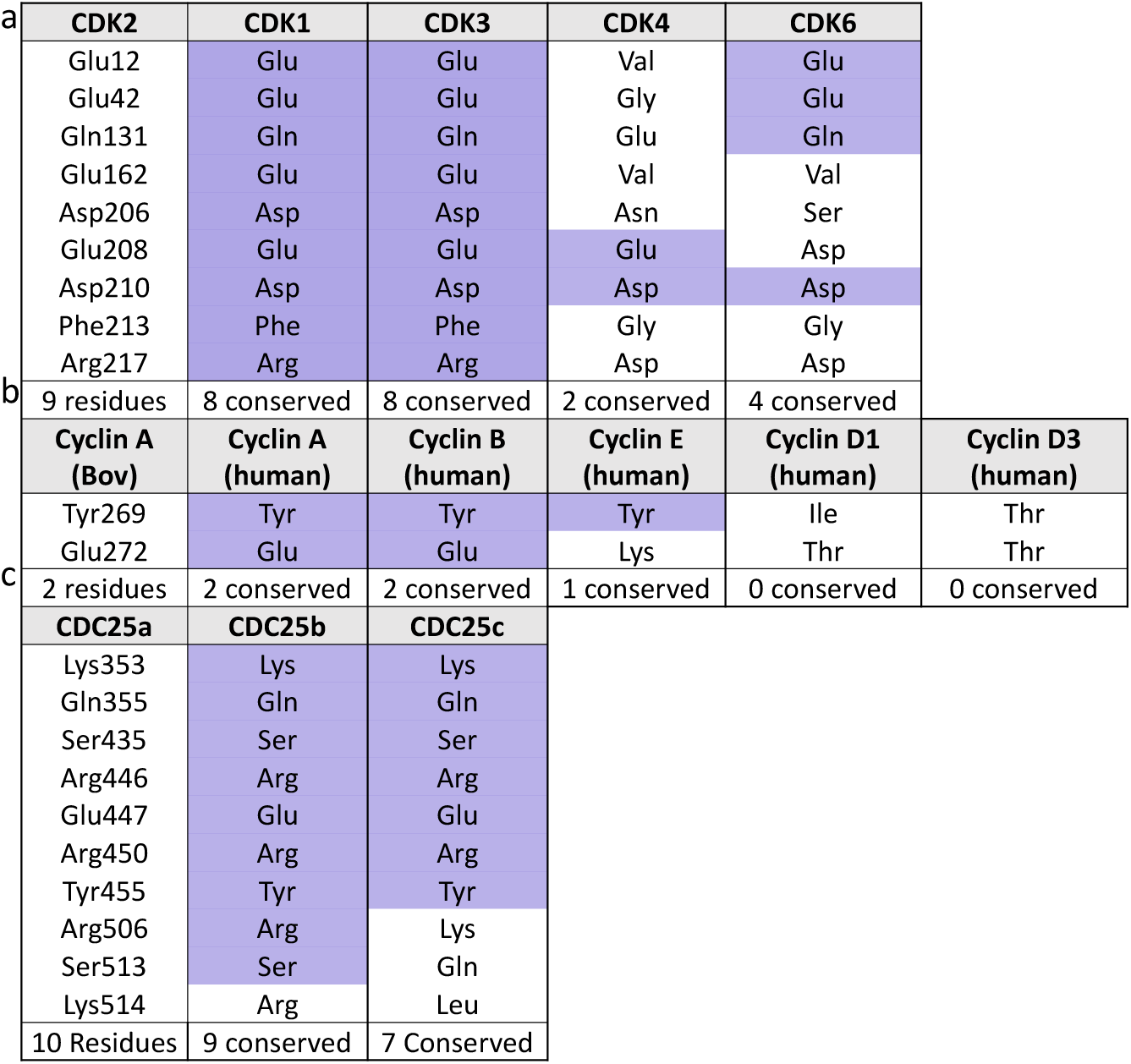
Sequence conservation of selected residues that mediate trimeric complex formation.

Selected residues identified from the cryo-EM structure that mediate complex formation. Conserved residues are indicated in purple. Complete sequence alignments are provided in Supplementary Figure 7.

Sequence alignment of CDK2 with CDKs 1, 3, 4 and 6 (Table. 1a, Supplementary Figure 7a-b) revealed that CDK1 and CDK3 are absolutely conserved at all 9 residues that mediate the CDK2-CDC25A interaction. In contrast, only 2 and 4 residues are conserved in CDK4 and CDK6 respectively (Table 1). Both CDK4 and CDK6 lack the GDSEID sequence (GNSEAD in CDK4 and GSSDVD in CDK6) and as the most prominent C-terminal interaction site, the sequence changes of CDK2 Asp206, Phe213 and Arg217 to asparagine, glycine and aspartic acid in CDK4 and to serine, glycine and aspartic acid in CDK6 would be expected to significantly impact their interaction with CDC25A (Fig. 7a-b).

**Figure 7:**
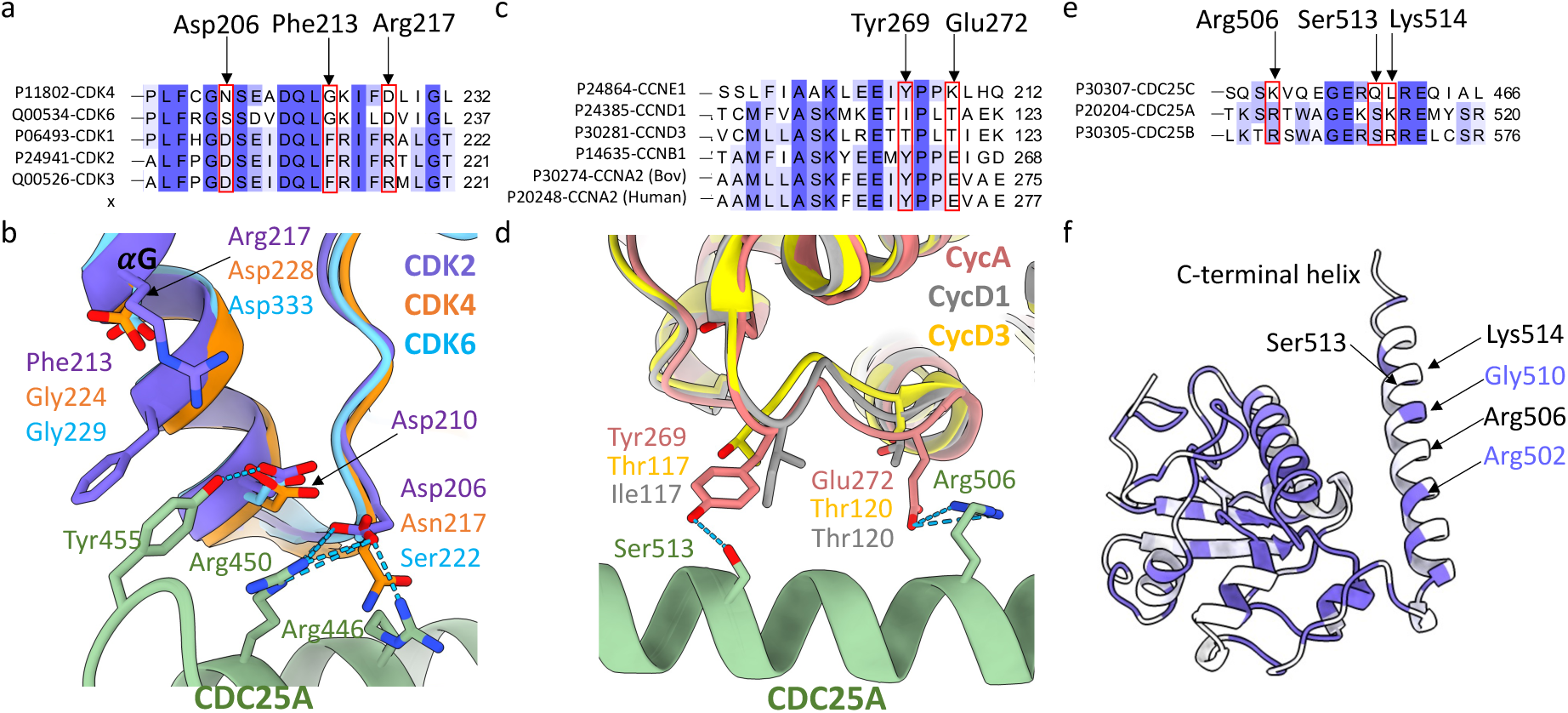
Sequence conservation at CDC25A binding regions. Sequence conservation is indicated in purple (darker shade = identical, lighter shade = similar, white = different) and selected residues directly involved in CDC25A binding are highlighted in red boxes. (a) Sequence conservation of CDK2 vs CDK1, 3, 4 and 6 around the GDSEID-αG helix motif. (b) Structural overlay of CDK2 (purple) with CDK4 (orange) and CDK6 (blue) showing changes in residue identity for Asp206, Phe213 and Arg217 at the GDSEID-αG helix motif (c) Sequence conservation of cyclin A vs cyclin B, E, D1 and D3 around the central helix (α3) in the cyclin A C-CBF. (d) Structural overlay of cyclin A (salmon) with cyclin D1 (grey) and cyclin D3 (yellow), highlighting the changes in residues identity for Tyr269 and Gly272 in the C-CBF. (e) Sequence conservation of CDC25A vs CDC25 B and C at the C-terminal helix region. (f) Ribbon diagram of CDC25A coloured by sequence conservation. CDC25A residues Arg506, Ser513 and Lys514 highlighted on the C-terminal helix are non-conserved in CDC25C.

Prior to the activation segment, CDK2 Gln131 is required to stabilise the extended CDC25A loop that binds in the wide groove between the N- and C-terminal lobes of CDK2. This residue is conserved in CDK6 but not in CDK4, in which Gln131 is mutated to glutamate. Within the activation segment, Glu162 hydrogen bonds to the CDC25A C-terminal helix but the inclusion of a valine at this position in the CDK4 and CDK6 sequences precludes this interaction. CDK4 and CDK6 also differ in sequence at key interacting residues within the N-terminal lobe. Whereas CDK6 conserves CDK2 N-terminal residues Glu12 and Glu42 (required to bind to the CDC25A active site and C-terminal helix respectively) they are mutated to valine and glycine respectively in CDK4.

Conservation analysis of cyclin A against B, E, and D-type cyclins (Table. 1b, Supplementary Figure 7c-d) is somewhat simplified given the relatively limited binding profile observed between CDC25A and cyclin A (Fig. 5, Table 1). Bovine cyclin A2 Tyr269 is conserved in cyclins B and E, but as a threonine in the D-type cyclins its ability to hydrogen bond to Ser513 on the CDC25A C-terminal helix is lost (Fig. 7c-d). Cyclin A Glu272, which forms a salt bridge with Arg506 on the CDC25A C-terminal helix, is also conserved in cyclin B, but changes to a lysine in cyclin E and to an isoleucine and threonine in cyclins D1 and D3 respectively. These residue changes in the D-type cyclins indicate binding of CDC25A may be compromised.

Ten CDC25A residues make significant interactions with CDK2-cyclin A, of which 9 and 7 are conserved in CDC25B and CDC25C respectively (Table. 1c, Supplementary Figure7e-f). CDC25A Arg446, Glu447, Arg450 and Tyr455 are key in promoting interaction with the CDK2 GDSEID motif and αG helix (Fig. 3i); these residues are conserved across CDC25B and C, as are CDC25A Ser435, Arg437 and Lys353 which bind the CDK2 G-loop (Fig. 3c). However, the CDC25A C-terminal helix residues that are vital for mediating interaction with both CDK2 and cyclin A (Fig. 5) have conservative changes in CDC25B; Arg506 and Ser513 are conserved and Lys514 changes to an arginine, but are non-conserved in CDC25C. Specifically, they are mutated to Lys452, Gln459 and Leu460 respectively (Fig. 7e-f). Combined these residue changes could significantly impact the engagement of the C-terminal helix of CDC25C to CDK2-cyclin A.

This sequence conservation analysis suggests that CDK1/2-cyclin A, CDK1-cyclin B and CDK2/3-cyclin E complexes are suitable binding partners for CDC25A. This is supportive of the notion that CDC25A aids in both G1/S and G2/M transitions. However, CDK4/6-cyclin D complexes appear unlikely substrates for CDC25A and, where they occur, may be dephosphorylated by other means. The detection of interactions between CDK4/6-cyclin D and CDC25 may result from CDK4/6 recognizing CDC25 proteins as substrates mediated through recruitment motifs within the CDC25 N-terminal sequence^39^.

As key regulators of cell cycle transitions and checkpoint pathways, CDC25A and CDC25B are frequently overexpressed in human cancers and often associated with poor prognosis. Consequently, the CDC25 isoforms are attractive yet challenging anticancer drug targets. Previous docking and molecular dynamic simulations have proved useful in predicting the overall binding mode of CDC25B^68^, and a fragment-based study has provided proof-of-concept in targeting the CDC25-CDK interaction^69^. We envisage this cryo-EM structure will facilitate the identification of allosteric sites that may be amenable to the development of inhibitors that prevent complex assembly or disrupt CDC25A catalysis.

## Methods

### Protein production, purification and complex formation

Human CDK2 phosphorylated on Tyr15 and Thr160, or only on Thr160, bovine cyclin A2, and human CDC25A and CDC25A mutant proteins were all expressed in recombinant *E. coli* cells and subsequently purified by affinity, ion-exchange and size exclusion chromatography steps. Detailed protocols for the various protein purifications are provided in the Supplementary Information.

#### Formation of the CDK2-cyclin A-CDC25A complex

To generate the trimeric complex, purified pY15pT160CDK2, cyclin A and CDC25A were mixed in a molar ratio of 1:1.5:2 and incubated on ice for 1hr. The complex was separated from monomeric and dimeric species by analytical gel filtration using an Akta Pure Microsystem (Cytiva) with a Superdex S200 10/300 column equilibrated in HBS (50 mM HEPES, 200 mM NaCl, 1 mM DTT, pH 7.4). Fractions containing the trimeric complex, as confirmed by SDS-PAGE, eluted from the column at a concentration suitable for cryo-EM studies (∼ 0.5 – 2.0 mg mL^-1^) without the need for concentration. The complex was snap-frozen in liquid nitrogen before application to cryo-EM grids.

### Cryo-EM Studies

All cryo-EM data were processed in CryoSPARC^70^ unless otherwise stated.

#### Initial sample preparation, data collection and processing

Preliminary cryo-EM samples were prepared using an FEI Vitribot mark IV set at 80 % humidity and 4 °C. 2.5 uL aliquots of the trimeric complex at ∼ 0.3 mg mL^-1^ were applied to QuantiFoil R2/2 200 mesh grids that had been glow discharged using a PELCO easiGlow™ Glow Discharge System for 1 min at 20 mAmp/0.26 mBar using ambient atmosphere. Grids were blotted with a blot force −10, blot time of 2-6 s and wait time of 3 s before plunge-freezing in liquid ethane.

Grids were screened and initial data collected on a 200 kV Glacios microscope (Thermo Fisher Scientific) with a Falcon 4 direct electron detector, housed at the York Structural Biology Laboratory (University of York, UK). Cryo-EM images were acquired at an accelerating voltage of 200 kV and nominal magnification of x240,000, giving a calibrated pixel size of 0.574 Å. Data were collected using AFIS in EER format, with an exposure time of 6.5 s and total fluence of 50 e/ Å^2^. A defocus range of −0.6 to −2.0 microns every 0.2 microns was used and autofocus was performed after centring. A 100micron objective aperture was used. A total of 11,955 movies were collected across two grids.

Movies were motion corrected using Patch Motion Correct Multi and the contrast transfer function (CTF) parameters were estimated using Patch CTF Correction Multi. Particles were picked using blob picker with a 40 Å minimum and 120 Å maximum particle diameter. Particle picks were inspected and filtered according to their power and NCC scores. Accepted particles were extracted with a box size of 384 pixels (px) and Fourier cropped to 96 px. Extracted particles were subject to multiple rounds of 2D classification using 40 iterations, a batch size of 200, and an initial classification uncertainty factor of 5. Significant preferential orientation was evident from the 2D classes. These particles were used to train Topaz picking (using the ResNet8 model architecture on a subset of 1,000 micrographs^52, 53^) and subsequent particle picking with the Topaz model provided additional 2D-views. Ab-initio reconstruction followed by homogenous refinement resulted in a map that encompassed the full complex, however, significant preferential orientation limited 3D refinement, leading to streaking and poor fitting to secondary structure.

#### Improving sample orientation distribution with CHAPS

To reduce the preferential orientation, samples were spiked with 0.1, 0.2, 0.5 and 1.0 X CMC (critical micelle concentration, CMC = 8 mM) CHAPS prepared in HBS buffer (50 mM HEPES, 200 mM NaCl, 1 mM DTT, pH 7.4) immediately prior to grid preparation. 2.5 uL aliquots of the trimeric complex (∼ 1.8 mg/mL) freshly spiked with CHAPS were applied to QuantiFoil R1.2/1.3 200 mesh grids that had been glow discharged using a PELCO easiGlow™

Glow Discharge System for 1 min at 20 mAmp/0.26 mBar using ambient atmosphere. The grids were blotted with a blot force −10, blot time of 4-12 s and wait time of 3s before plunge-freezing in liquid ethane using a FEI Vitribot mark IV set at 80% humidity and 4 °C.

Grids were screened, and initial data were collected on a Glacios microscope (Thermo Fisher Scientific) with a Falcon 4 direct electron detector (York Structural Biology Laboratory). At each CHAPS concentration, ∼ 1000 exposures were collected at an accelerating voltage of 200 kV, nominal magnification x240,000, a calibrated pixel size of 0.574 Å, exposure time of 3.2 s and a total fluence of 50 e/Å^2^. Data collection was set up using the multigrid function in the EPU Software (Thermo Fisher Scientific) as described above for the initial data collection. All data sets were processed separately to 2D classification to identify the CHAPS concentration most beneficial for improving the particle orientation distribution. Movies were motion corrected using Patch Motion Correct Multi and the contrast transfer function (CTF) parameters estimated using Patch CTF Correction multi. Particles were picked using blob picker with a 40 Å minimum and 120 Å maximum particle diameter. Particle picks were inspected, filtered according to their power and NCC scores and extracted with a box size of 384 px, Fourier cropped to 96 px to give a pixel size of 2.296 Å. Extracted particles were subject to multiple rounds of 2D classification using 40 iterations, a batch size of 200 and an initial classification uncertainty factor of 5. The positive effect of CHAPS on the particle orientation distribution was evident at all concentrations tested, however, more unique views were identified at 0.5 and 1.0 X CMC. These grids were chosen for further data collection.

#### Final high resolution data collection

The final high resolution data set was collected across 2 grids (Quantifoil 1.2/1.3 200 mesh) prepared with 1.8 mg mL^-1^ trimeric complex and 0.5 and 0.1 X CMC CHAPS. Cryo-EM images were acquired on a 300 kV FEI Titan Krios equipped with a BioQuantum K3 detector and Gatan Imaging Filter (GIF), housed at the Electron Bio-Imaging Centre (Krios 1, eBIC, Diamond Light Source, UK). Data were collected at an accelerating voltage of 300 kV and x150,000 magnification, yielding a pixel size of 0.825 Å, in counting mode with an energy filter slit width set at 20 eV. Data collection was set up using the multigrid function in the EPU Software (Thermo Fisher Scientific) using AFIS with 3 exposures per hole. Data were collected in fifty frame movies in TIFF format, with an exposure time of 1.92 s, a dose of 1.010 e/Å^2^ per frame and a total accumulative dose of 50.49 e/Å^2^. A defocus range of −0.6 to −2.0 microns every 0.2 microns was used and autofocus was performed after centring. A 100-micron objective aperture was used. A total of 13,242 movies were recorded.

#### Data processing, and structure determination

Movies were patch motion corrected using Patch Motion Correct Multi and the contrast transfer function (CTF) parameters estimated using Patch CTF Correction multi. Particles were picked using automated blob picker with a 40 Å minimum and 120 Å maximum particle diameter, resulting in ∼ 6,000,000 particle picks. Particles were inspected and filtered according to their power and NCC scores to yield a particle set of 4,981,705 particles. Accepted particles were extracted with a box size of 288 px, Fourier cropped to 96 px to give an effective pixel size of 2.475 Å. Extracted particles were subjected to multiple rounds of 2D classification using 40 iterations, a batch size of 200 and an initial classification uncertainty factor of 5 (same parameters applied to all 2D classification steps unless otherwise specified). Following 2D classification, 1,231,048 particles from 76 classes were used to generate 2 ab-initio models, with a 0.01 class similarity cut-off. Class 0 of 757,792 particles was chosen as the best initial model, and particles were re-extracted with a box size of 288 px without Fourier cropping to give a pixel size of 0.825 Å. Using class 0 ab-initio as the starting model, 3D refinement was performed using the homogeneous refinement with a 10 Å maximum alignment resolution, followed by non-uniform refinement^71^ (no max align) to yield a 3.2 Å (FSC 0.143) cryo-EM map of the trimeric complex.

To remove further heterogeneity, particles constituting this reconstruction were classified by 3D heterogenous refinement using the previously generated ab-initio models and a refinement box size of 144 px. Class 0 (534,740 particles, 5.6 Å) was selected as the best class, re-extracted with a box size of 288 px and refined by homogenous refinement (10 Å maximum alignment resolution) and non-uniform refinement to generate an improved reconstruction at 3.05 Å resolution (FSC 0.143).

Particles constituting this reconstruction were used to train Topaz using the ResNet8 model architecture on a subset of 1,000 micrographs previously denoised with Topaz Denoise^52, 53^. The resulting Topaz model was used for picking from the full set of 13,242 Topaz denoised micrographs. Following Topaz picking, 2,105,240 particles were extracted with a box size of 288 px, cropped to 144 px with an effective pixel size of 1.65 Å, and 2D classified to remove poor particles. The remaining 1,898,974 particles were sorted into 6 ab-initio models using a 0.01 class similarity cut-off, and further 3D classified and refined by heterogeneous refinement using the 6 ab-initio models. Particles from the best 3D heterogenous class (670,852 particles) were re-extracted without Fourier cropping to give an effective pixel size of 0.825 Å and refined by homogeneous with a 10 Å maximum alignment resolution. The 3D reconstruction was further refined by non-uniform refinement^71^ to yield the final 2.91 Å reconstruction of the trimeric complex. The local resolution was estimated via the local resolution estimation job in CryoSPARC (0.143 FSC threshold).

#### Model generation, refinement and validation

The resolution of the 3D reconstruction was sufficient to permit automated model building using ModelAngelo^54^, which was performed using a FASTA sequence containing the expressed residues of all three proteins. The model generated by ModelAngelo showed good fitting to the EM map, however, the first 26 N-terminal residues of CDK2 (containing the G-loop) and two loop regions of CDC25A (413-424 and 151-154) were modelled with low confidence. These regions could be discerned by manual modelling according to the EM map, to generate a complete model of the trimeric complex.

The completed model and EM map were imported into Phenix^56^ for model refinement and validation. The fitting of the model was improved with real-space refinement followed by manual refinement in COOT^55^ based on MolProbity^72^ outlines. The model, cryo-EM map and map-to-model quality was assessed in Phenix using Comprehensive Validation^73^ before deposition to the Protein Data Bank (PDB **8QKQ**) and the Electron Microscopy Data Bank (EMD-**18470**) Full data collection, refinement and validation statistics are given in Supplementary Table. 1.

All structural figures were generated in UCSF ChimeraX^74, 75^.

#### Homogenous time resolved fluorescence

HTRF assays were carried out essentially as described^76^. GST-CDC25A wild-type and mutant proteins (at 100 nM) and biotinylated Avi-tagged pT160CDK2-cyclin A (at an 11-point 2-fold dilution series starting at 8 μM) were prepared in HTRF buffer A (50 mM HEPES, 100 mM NaCl, 1 mM DTT and 0.1 mg mL^-1^ BSA) and incubated together for 60 min at 4°C. 4 nM Tb labelled anti-GST antibody and SAXL665 at 1/8th the concentration of the biotinylated pT160CDK2-cyclin A, were prepared in HTRF buffer B (50 mM HEPES, 100 mM NaCl, 1mM DTT and 0.1 mg mL^-1^ BSA) and added to each well. The plate was incubated for a further 120 min at 4°C, before being scanned. Samples were excited using a wavelength of 337 nm and emission spectra measured at 620 nm and 665 nm (PHERAstar FS (BMG LABTECH)). Binding curves were plotted using GraphPad Prism 6 from which the K_d_s were determined. The curves shown are representative binding curves from at least two biological replicates each run in triplicate and carried out on separate days.

#### Hydrogen-deuterium exchange mass spectrometry

HDX-MS experiments were carried out using an automated HDX robot (LEAP Technologies, Fort Lauderdale, FL, USA) coupled to an M-Class Acquity LC and HDX manager (Waters Ltd., Wilmslow, Manchester, UK). Proteins (pY15/pT160CDK2, cyclin A, CDC25A, pY15/pT160CDK2-cyclin A and CDK2-cyclin A-CDC25A) were diluted to 10 µM in equilibration buffer (40 mM HEPES pH 7.4 150 mM NaCl, 1mM TCEP.HCl) prior to analysis. 5 µl sample was added to 95 µl deuterated buffer (40 mM HEPES pD 7.4 150 mM NaCl, 1mM TCEP.HCl) and incubated at 4 °C for 0.5, 2, 10 or 30 min with each condition being acquired in triplicate along with three t=0 conditions using protonated buffer to provide the baseline mass. Following the labelling reaction, samples were quenched by adding 75 µl of the labelled solution to 75 µl quench buffer (50 mM potassium phosphate, 0.05% DDM which was pH adjusted to give a final quenched pH of ∼ 2.5). 50 µl of quenched sample were passed through a home-packed pepsin column using agarose immobilised pepsin (Thermo Fisher Scientific) at 40 µl min^−1^ (20 °C) and a VanGuard Pre-column Acquity UPLC BEH C18 (1.7 µm, 2.1 mm × 5 mm, Waters Ltd., Wilmslow, Manchester, UK) for 3 min in 0.3% formic acid in water. The resulting peptic peptides were transferred to a C18 column (75 µm × 150 mm, Waters Ltd., Wilmslow, Manchester, UK) and separated by gradient elution of 0–40% MeCN (0.1% v/v formic acid) in H2O (0.3% v/v formic acid) over 12 min at 40 µl min^−1^. Trapping and gradient elution of peptides was performed at 0 °C. The HDX system was interfaced to a Synapt G2Si mass spectrometer (Waters Ltd., Wilmslow, Manchester, UK) using electrospray ionisation with a capillary voltage of 3 kV, cone gas of 50 L/h, desolvation gas 300 L/h and nebuliser of 6.5 bar. Desolvation and source temperatures were 120 and 80 °C respectively. HDMSE, dynamic range extension and sensitivity modes (Data Independent Analysis (DIA) coupled with IMS separation and single reflectron time-of-flight) were used to separate peptides prior to CID fragmentation in the transfer cell. Argon pressure in the trap and transfer cells was 2.4×10^-2^ mbar, helium pressure was 4.21 mbar and nitrogen pressure in the drift tube was 3 mbar. Wave heights and velocities were 311 m/s and 4 V for the trap, 650 m/s and 30 V for the IMS and 175 m/s and 4V for the transfer cell. Signals were acquired over m/z 50-2000 for 0.6 s alternating between high and low energy scans with a collision energy ramp of 18-40 V in the high energy scan to induce peptide fragmentation. HDX data were analyzed using PLGS (v3.0.2) and DynamX (v3.0.0) software supplied with the mass spectrometer. A minimal database was used for peptide ID containing the target sequences and pepsin. Restrictions for identified peptides in DynamX were as follows: minimum intensity: 2500, minimum products per MS/MS spectrum: 3, minimum products per amino acid: 0.3, maximum sequence length: 18, maximum ppm error: 10, file threshold: 4/6. Following manual curation of the data, hybrid plots were generated using Deuteros 2.0 at 0.01 significance^77^.

## Abbreviations

CDK: cyclin-dependent kinase
CBF: cyclin-box fold
SEC: size exclusion chromatography
HDX: Hydrogen-Deuterium Exchange
HTRF: Homogenous Time-Resolved Florescence
FSC: Fourier shell correlation
CTF: contrast transfer function.

## Data availability

Data are available upon request to the corresponding author. The map and model of the trimeric CDK2-cyclin A-CDC25A complex has been deposited in the Protein Data Bank and the Electron Microscopy Data Bank with accession numbers PDB **8QKQ** and EMD-**18470.**

## Acknowledgements

This research was supported by the Medical Research Council (Grant References MR/N009738/1 and MR/V029142/1). Cryogenic electron microscopy was carried out at the Electron Bio-imaging Centre (eBIC) at the Diamond Light Source under the Northern England Cryo-EM Consortium (proposal BI28576-22), and the Cryo-EM facility at the University of York (Glacios microscope funded by The Wellcome Trust grant number 206161/Z/17/Z). We wish to thank Dr Davide Zabeo at the Electron Bio-imaging Centre for facilitating Cryo-EM data collections and Dr Arnaud Baslé at Newcastle University for assistance with data management. We thank colleagues who contributed to the early stages of this work; T. Johnson, M. Morgan, S. Platsaki, W. Stanley and K. Yata. The LEAP sample handling robot used in the HDX-MS work was a kind donation from Waters UK. For the purpose of open access, the authors have applied a Creative Commons Attribution (CC BY) licence to any Author Accepted Manuscript version arising from this submission.

## Author Contributions

Rhianna Rowland: Conceptualization, methodology (cryo-EM data acquisition), investigation (protein production, cryo-EM analysis and structure determination), visualisation and writing (original draft, review and editing). Svitlana Korolchuk: methodology (protein production) and investigation (HTRF assays). Marco Salamina: Conceptualization, investigation, collaboration. James Ault: Investigation (HDX data collection and analysis).

Sam Hart: Resources (facilitation of cryo-EM data acquisition. Johan Turkenburg: Resources (facilitation of cryo-EM data acquisition). James Blaza: Funding acquisition, cryo-EM data acquisition. Jane Endicott: Conceptualization, resources, supervision, funding acquisition, writing (original draft, review and editing). Martin Noble: Conceptualization, resources, supervision, funding acquisition, writing (original draft, review and editing). All authors reviewed the manuscript.

## Competing Interests

The other authors declare no competing interests. The authors declare no competing financial interest. Some work in the authors’ laboratory is supported by a research grant from Astex Pharmaceuticals. SK is currently an employee of FujiFilm, Billingham, UK; MS is currently and employee of Evotec, Milton, UK.

## Additional information

**Correspondence** and requests for materials should be addressed to Jane A. Endicott or Martin E.M. Noble.

