## Supplementary Material for "Cryo-EM structure of the CDK2-cyclin A-CDC25A Complex"

**Inventory of Supplemental Information**

**Supplementary Methods**

**Supplementary Figures**
Supplementary Figure 1, associated with Figure 1
Supplementary Figure 2, associated with Figure 1
Supplementary Figure 3, associated with Figure 1
Supplementary Figure 4, associated with Figure 1
Supplementary Figure 5, associated with Figure 2
Supplementary Figure 6, associated with Figure 5
Supplementary Figure 7, associated with Table 1

**Supplementary Tables**
Supplementary Table 1

**Supplementary References**

**Supplementary Methods**

*Production of pY15pT160CDK2*

GST-3C tagged human CDK2 (Uniprot 24941, residues 1-298) and GST-tagged kinase domain of human Wee1 were cloned into a single pGEX6P1 vector to facilitate Tyr15 phosphorylation of CDK2. This construct yields the pY15CDK2 protein sequence preceded by Pro-Gly-Ser as a cloning artefact following 3C cleavage. The CDK2-Wee1 co-expression vector was transformed in BL21(DE3)STAR cells and grown at 37 °C to OD ~ 0.8 before reducing the temperature to 18 °C and inducing with 0.1 mM IPTG. Cells were incubated for ~24 hr and harvested by centrifugation (4,000 xg, 20 min, 4 °C). The cell pellet was resuspended in modified HBS buffer (40 mM HEPES, 200 mM NaCl, pH 7.4, 1 x-EDTA tablet/50 mls, 1 mM DTT, 2 mg $mL^{-1}$ DNase I, 10 mg $mL^{-1}$ RNase A, 25 mg $mL^{-1}$ lysozyme and 5 mM $MgCl_2$) and lysed via sonication (5 min total, pulsed 20 s on and 40 s off, 30 % amp on ice) before clarifying by centrifugation (60,000 xg, 60 min, 4 °C).

All purification steps were performed at 4°C. The clarified lysate was bound to glutathione sepharose 4B resin (Cytiva), and pre-equilibrated in HBS (40 mM HEPES, 200 mM NaCl, 2 mM DTT, pH 7.4) using a gravity flow column. Protein was eluted with 20 mM reduced glutathione in HBS pH 7.4, then cleaved overnight (16 hr) at 4 °C using 1:50 w/w 3C protease:GST-CDK2. Cleaved product was further purified by size exclusion chromatography (SEC) on an S75 16/600 Superdex® column (Cytiva) equilibrated in HBS pH 7.4 with 2 mM DTT to yield purified pY15CDK2.

To prepare doubly phosphorylated pY15pT160CDK2, monomeric pY15CDK2 was phosphorylated by *S. cerevisiae* CAK1 *in vitro*. We note that *S. cerevisiae* CAK1 efficiently phosphorylates monomeric CDK2 *in vitro* but not the cognate cyclin-bound complexes, necessitating individual expression of each complex component. GST-CAK1 cloned into a pACEBac1 vector, was expressed and purified from recombinant insect cells and purified by a single affinity chromatography step with glutathione sepharose 4B resin (GE Cytiva) as described above. Purified CDK2 and GST-CAK1 were mixed in a 4:1 molar ratio CDK2:CAK1 in HBS buffer pH7.5 supplemented with 1 mM ATP, 100 mM $MgCl_2$, 50 mM Tris-HCl and phosphatase inhibitor, and incubated overnight at 4 °C. The extent of CDK2 phosphorylation was determined by intact mass spectrometry analysis. GST-CAK1 was separated from pY15pT160CDK2 by subtractive GST-affinity purification on glutathione sepharose 4B resin

(Cytiva) in gravity flow format. pY15pT160CDK2 was further purified by analytical SEC on a Superdex 200 10/300 column equilibrated in HBS pH7.5 supplemented with 1 mM DTT on an Akta Pure Micro system (Cytiva). Fractions containing pY15pT160CDK2 were confirmed by SDS-PAGE, pooled and concentrated to 1.5 mg $mL^{-1}$ using a Vivaspin 10 kDa MWCO centrifugal filter (Sartorius).

*Production of bovine cyclin A*

GST-3C tagged bovine cyclin A2 (UniProt P30274, residues 169-430) was cloned into a pGEX6P1 vector to yield the cyclin A sequence preceded by Pro-Leu-Gly-Ser-Met-Gly as a cloning artefact following 3C cleavage. Bovine cyclin A2 was used as the equivalent human cyclin A2 construct is prone to aggregation, and assembly of the complex necessitated each component being expressed and purified individually. This construct was transformed into BL21(DE3) Rosetta cells which were grown at 37 °C to an OD ~ 0.4-0.6, before dropping the temperature to 18 °C and inducing with 0.1 mM IPTG. Cells were incubated at 18 °C overnight and harvested by centrifugation (4,000 xg, 20 min, 4 °C). Cell pellets were resuspended in 50 mM Tris, 300 mM NaCl, 100 mM $MgCl_2$ pH 8.0, 1 x-EDTA tablet /50 mls, 1 mM DTT, 2 mg $mL^{-1}$ DNase I, 10 mg $mL^{-1}$ RNase A, 25 mg $mL^{-1}$ lysozyme and 5 mM $MgCl_2$. Cell suspensions were lysed via sonication (5 min total, pulsed 20 s on and 40 s off, 30 % amp on ice) and clarified by centrifugation (60,000 xg, 60 min, 4 °C).

All purification steps were performed at 4°C. Clarified lysate was bound to glutathione sepharose 4B resin (Cytiva), pre-equilibrated in 50 mM Tris, 300 mM NaCl, 100 mM $MgCl_2$, pH 8.0 supplemented with 1 mM DTT, using a gravity flow column. Protein was eluted with 20 mM reduced glutathione in 50 mM Tris, 300 mM NaCl, 100 mM $MgCl_2$ pH 8.0 and cleaved overnight (16 hr) at 4 °C using 1:50 w/w 3C protease:GST-cyclin A. Cleaved product was further purified by SEC on S75 26/600 Superdex® HiLoad® column (Cytiva), equilibrated in 50 mM Tris, 300 mM NaCl, 100 mM $MgCl_2$, pH 8.0 supplemented with 1 mM DTT. Fractions containing cyclin A were identified by SDS-PAGE, pooled and concentrated to 3.4 mg $mL^{-1}$ using a Vivaspin 10 kDa MWCO centrifugal filter (Sartorius).

*Production of CDC25A*

The catalytic and C-terminal domain sequence of CDC25A (Uniprot entry P30304, residues 335-524) was cloned into a pGEX6P1 vector with a 3C cleavable GST tag, and made catalytically inactive by introducing the Cys430Ser mutation, to yield the inactive CDC25A sequence preceded by Pro-Leu-Gly-Ser as a cloning artefact following 3C cleavage. This construct was transformed into BL21(DE3) pLysS and grown at 37 °C to an $OD_{600\ nm}$ ~0.4-0.6, before dropping the temperature to 18 °C and inducing with 0.1 mM IPTG. Cells were incubated at 18 °C overnight and harvested by centrifugation (4,000 xg, 20 min, 4 °C) and resuspended in modified HBS buffer (40 mM HEPES, 200 mM NaCl, pH 7.4, 1 x-EDTA tablet /50 mls, 1 mM DTT, 2 mg $mL^{-1}$ DNase I, 10 mg $mL^{-1}$ RNase A, 25 mg $mL^{-1}$ lysozyme and 5 mM $MgCl_2$). Cell suspensions were lysed via sonication (5 min total, pulsed 20 s on and 40 s off, 30 % amp on ice) and clarified by centrifugation (60,000 xg, 60 min, 4 °C).

All purification steps were performed at 4°C. The clarified lysate was bound to glutathione sepharose 4B resin (Cytiva), pre-equilibrated in HBS (40 mM HEPES, 200 mM NaCl, pH 7.4) supplemented with 1 mM DTT, using a gravity flow column. Protein was eluted with 20 mM reduced glutathione in HBS pH 7.4, then cleaved overnight (16 hr) at 4 °C using 1:50 w/w of 3C protease: GST-CDC25A. Cleaved product was further purified by SEC on an S75 preparative 26/600 Superdex® HiLoad® column (Cytiva), equilibrated in HBS, supplemented with 2 mM DTT, pH 7.4. Fractions containing CDC25A were identified by SDS-PAGE and pooled. Contaminating DNA was removed by cation-exchange chromatography using a HiTrap-SP sepharose FF 5mL column (Cytiva) pre-equilibrated in 20 mM HEPES, 75 mM NaCl, pH 7.4 supplemented with 1 mM DTT. CDC25A was eluted via a linear gradient over 20 column volumes into 100% high salt buffer containing 20 mM HEPES, 1 M NaCl, pH 7.4, 1 mM DTT. Fractions containing CDC25A were identified by SDS-PAGE, concentrated to 2 mg $mL^{-1}$ using a Vivaspin 10 kDa MWCO centrifugal filter (Sartorius). For homogenous time-resolved fluorescence (HTRF) assays, GST-CDC25A was prepared as described above except that following the initial affinity column purification, the cleavage step was omitted, the eluate was concentrated and then further purified by SEC. GST-CDC25A mutants were generated by site-directed mutagenesis using the QuikChange method (Agilent Technologies), confirmed by sequencing (Eurofins), and then expressed and purified as described above.

*Production of biotinylated pT160CDK2-cyclin A*

N-terminally Avi-tagged human CDK2 phosphorylated at Thr160 was produced in *E. coli* by co-expression of human GSTAviCDK2 and *S. cerevisiae* GSTCAK1 from the pGEX6P-1 vector backbone (GE Healthcare). The introduction of the Avi-tag at the N-terminus of CDK2 generates the full-length CDK2 protein preceded by the sequence GPAM*GLNDIFEAQKIEWHE*A (Avi tag residues italicised). pT160CDK2 was then expressed as described above for the generation of monomeric pY15CDK2. Bovine cyclin A was expressed as described above. To generate the complex the pellets from cultures expressing CDK2 and cyclin A were thawed and mixed in a ratio of 1:2 by culture volume and supplemented by addition of 10 µg $mL^{-1}$ RNase A, 2 µg $mL^{-1}$ DNaseI, 25 µg $mL^{-1}$ lysozyme and 5 mM $MgCl_2$. The mixed cell suspension was lysed via sonication (5 min total, pulsed 20 s on, and 40 s off, 30 % amp on ice) and clarified by centrifugation (48,000 xg, 60 min, 4 °C). The supernatant was filtered through a 0.45 µm filter and applied to a glutathione sepharose 4B column (Cytiva) pre-equilibrated in 50 mM Tris pH 7.5, 150 mM NaCl, 0.05%TCEP (TBS) and subsequently eluted in the same buffer supplemented with 20 mM glutathione. The GST tag was cleaved by 3C protease (1:50 w/w ratio) incubated overnight at 4 $^{o}$C, and then subsequently further purified by size exclusion chromatography (Superdex 75 26/60 column (Cytiva) equilibrated in HBS). Fractions containing AvipT160CDK2-cyclin A were identified by SDS-PAGE and pooled. Co-eluting glutathione-S-transferase was removed by subjecting the sample to a subtractive glutathione-sepharose 4B column and collecting the flow-through. AvipT160CDK2-cyclin A was then concentrated to 40-50 µM and added to the biotinylation solution (50 mM Bicine buffer pH 8.3, 10 mM ATP, 10 mM Mg acetate, 100 µM $\alpha$-biotin, final concentrations supplemented with 75 µg BirA) and incubated with gentle rotation at 4 $^{o}$C overnight. The sample was then concentrated to 1 ml, desalted using a HiTrap 5ml prepacked desalting column (Cytiva) into HBS. Aliquots were flash frozen and stored at -80 $^{o}$C.

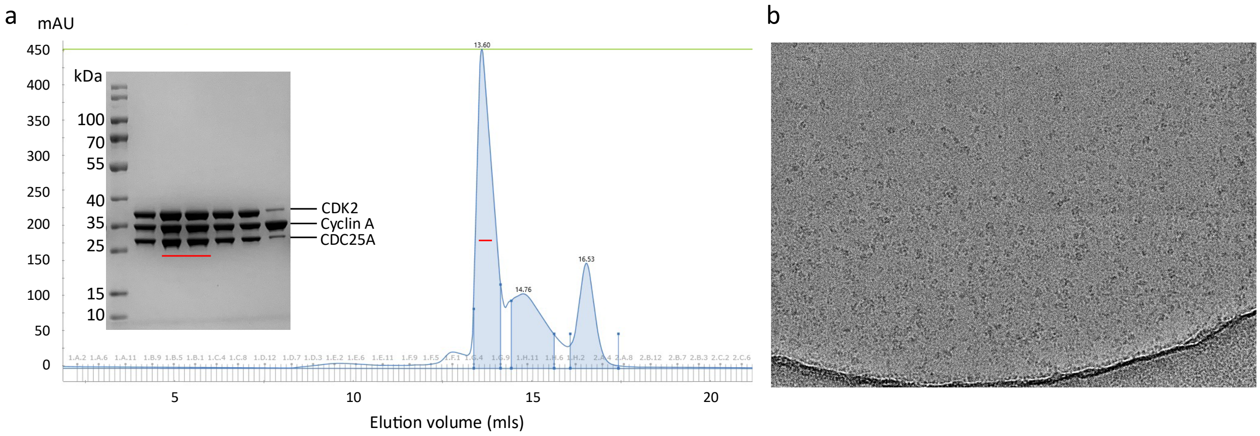

**Supplementary Figure 1: CDK2-cyclin A-CDC25A complex purification and cryo-EM sample preparation**

a) Following independent expression and purification, CDC25A was incubated with pY15pT160CDK2 and cyclin A to form the trimeric complex which was isolated by analytical gel filtration using an S200 10/300 column. Red bar on the SDS-PAGE gel (inset) and on the SEC chromatogram identifies the pY15pT160CDK2-cyclin A-CDC25A sample taken for cryo-EM analysis. (b) The specimen was supplemented with 0.5-1.0 X CMC CHAPS before application to Quanitfoil 1.2/1.3 holey carbon grids. Representative cryogenic electron micrograph showing CDK2-cyclin A-CDC25A particles imaged on a 300 kV Titan Krios with a K3 detector and GIF energy filter.

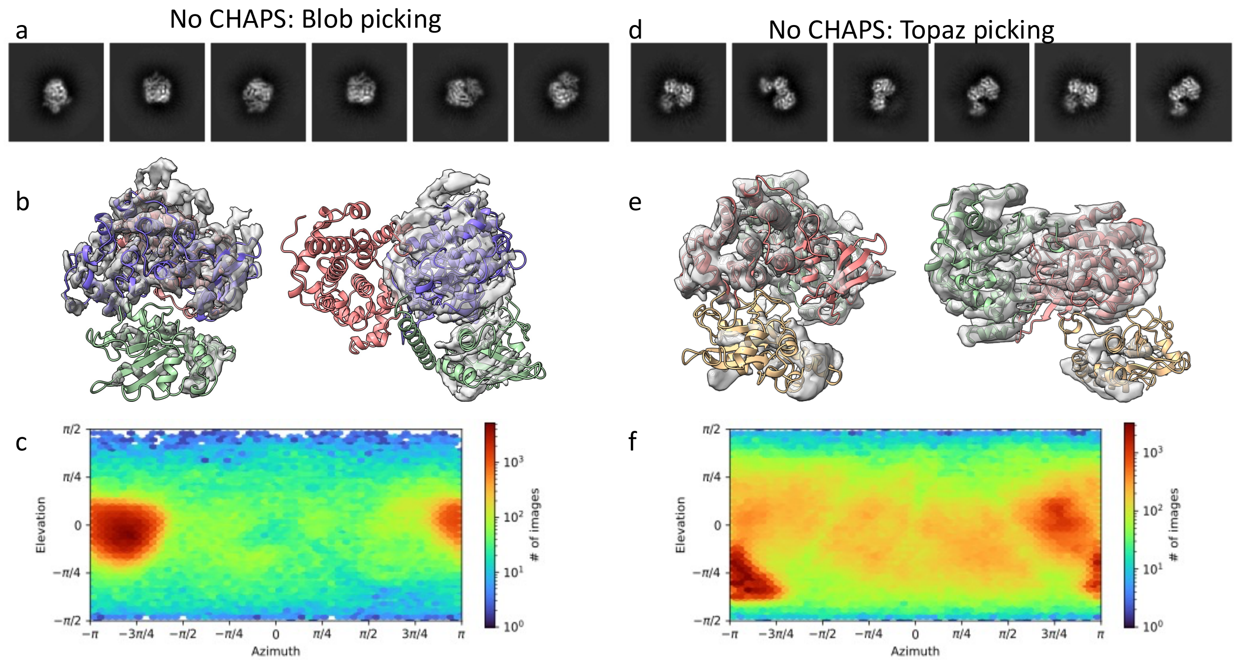

**Supplementary Figure 2: Preliminary cryo-EM data of the trimeric complex in the absence of CHAPS detergent**

(a-c) In the absence of CHAPS, blob picking of the trimeric complex yielded (a) 2D class averages of limited unique views; all classes represented a front view of the CDK2-CDC25A motifs, with no views for the cyclin A unit. (b) This significant preferential orientation affected 3D refinement; whilst EM density for the CDK2 (purple) and CDC25A (green) units was resolved, the map lacked density for cyclin A (salmon). (c) Preferential orientation is indicated by the orientation distribution heat map of the refined particle set. (d-f) Topaz picking of the trimeric complex improved the number of unique views (d) with a wider variety of 2D class averages produced. (e) This resulted in a 3D reconstruction that encompassed all three protein units (CDK2 red, cyclin A green and CDC25A yellow), however, preferential orientation persisted, and the reconstruction exhibited streaking in the EM density. (f) Improved orientation distribution heat map from Topaz picked particles.

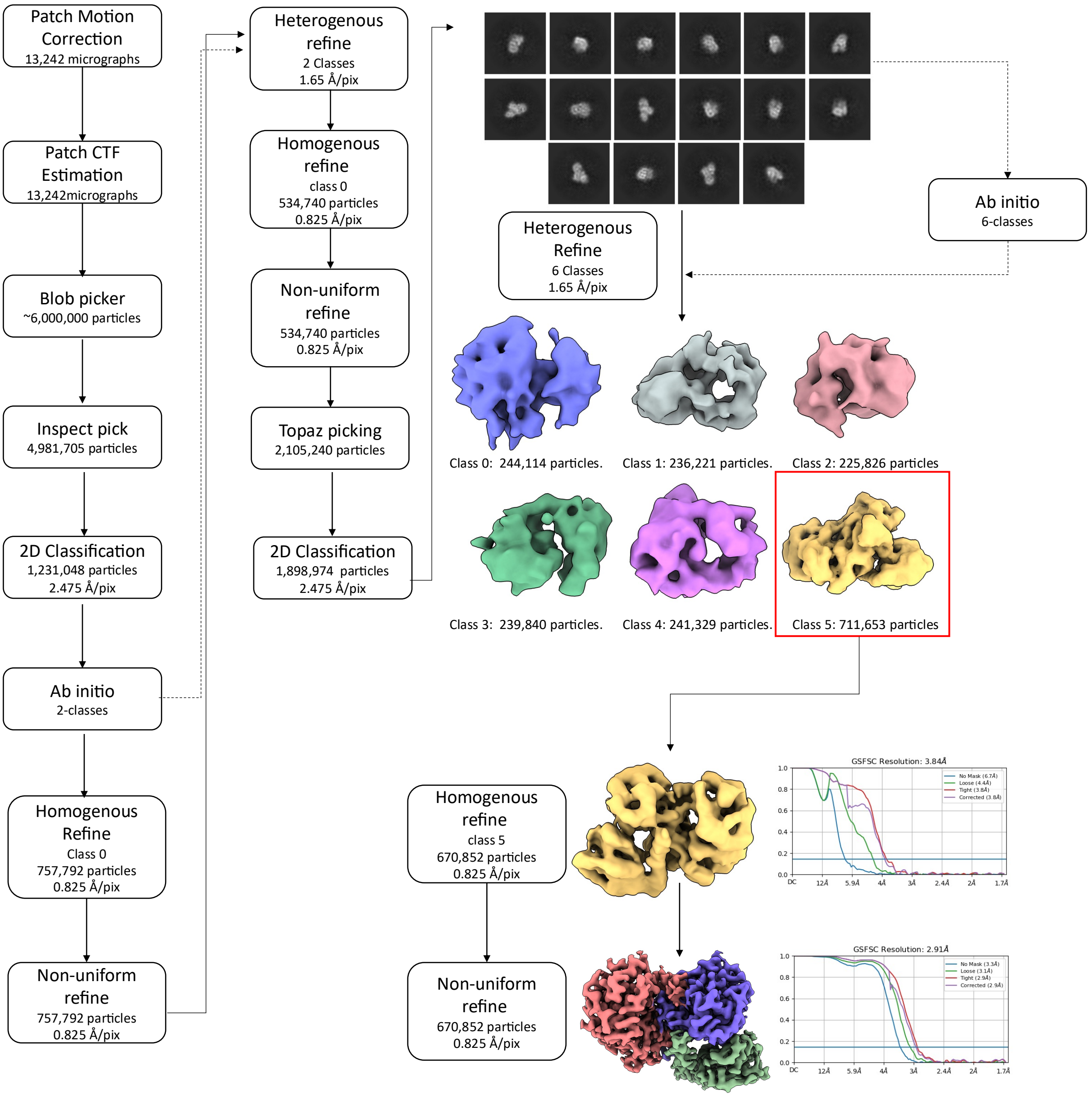

**Supplementary Figure 3: Structure determination of the CDK2-cyclin A-CDC25A complex by Cryo-EM**

Movies were patch motion corrected and the contrast transfer function (CTF) estimated. Using automated-blob picker, ~ 6,000,000 particles were picked and filtered according to their power and NCC scores to yield a particle set of 4,981,705 particles. Accepted particles were extracted with a pixel size of 2.475 Å and subjected to multiple rounds of 2D classification. The resulting 1,231,048 particles were used to generate 2 ab-initio models; class 0 was re-extracted with a pixel size of 0.825 Å and refined by homogeneous refinement and non-uniform refinement. 3D heterogenous refinement was performed using 2-classes, with class 0 (534,740 particles, 5.6 Å) selected as the best class and further refined by homogenous and non-uniform refinement. These particles were used to train Topaz using the ResNet8 model on a subset of 1,000 micrographs. Following Topaz picking, 2,105,240 particles were extracted with a pixel size of 1.65 Å and 2D classified to remove poor particles. The remaining 1,898,974 particles were sorted into 6 ab-initio models before further refinement by heterogenous refinement using the 6 ab-initio classes. Particles from class 5 (670,852 particles) were re-extracted with an effective pixel size of 0.825 Å before refinement by homogenous and non-uniform refinement to yield the final 2.91 Å reconstruction (FSC 0.143).

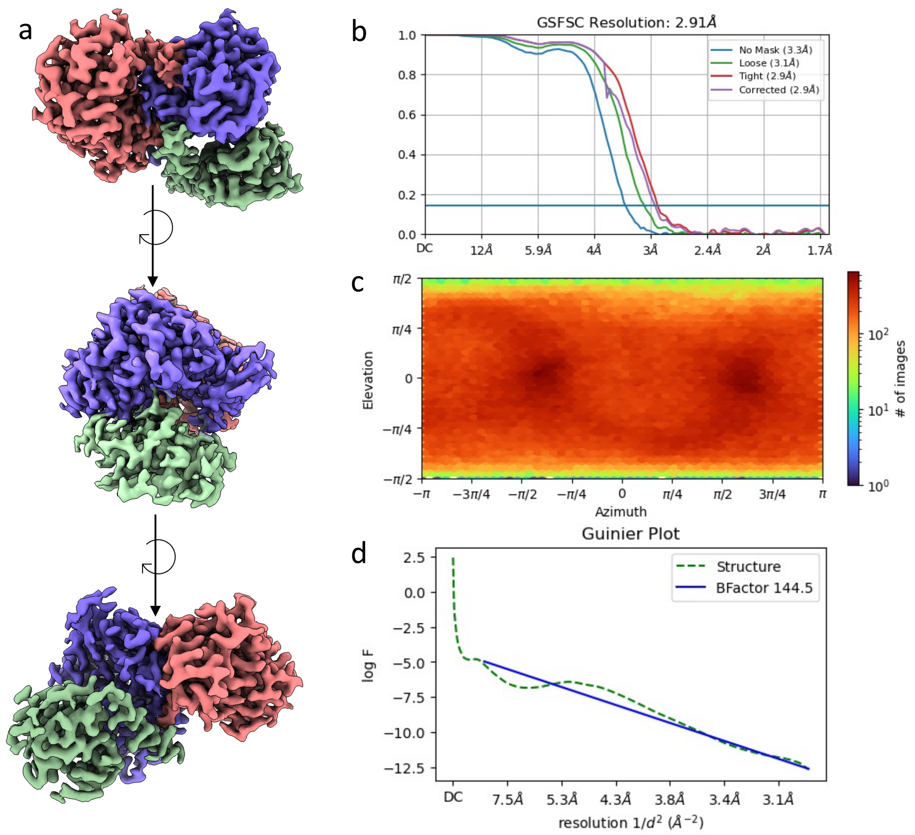

**Supplementary Figure 4: Cryo-EM map of the CDK2-cyclin A-CDC25a trimeric complex**

(a) Views of the trimeric reconstruction comprising cyclin A (salmon), CDK2 (purple) and CDC25A (green). (b) Gold standard Fourier shell correlation plot for the refined 2.91 Å reconstruction (FSC 0.143). (c) Orientation distribution heat map of the refined particle set constituting the final 3D reconstruction. (d) Guinier plots of final refinement; global B-factor = 144 Å$^2$.

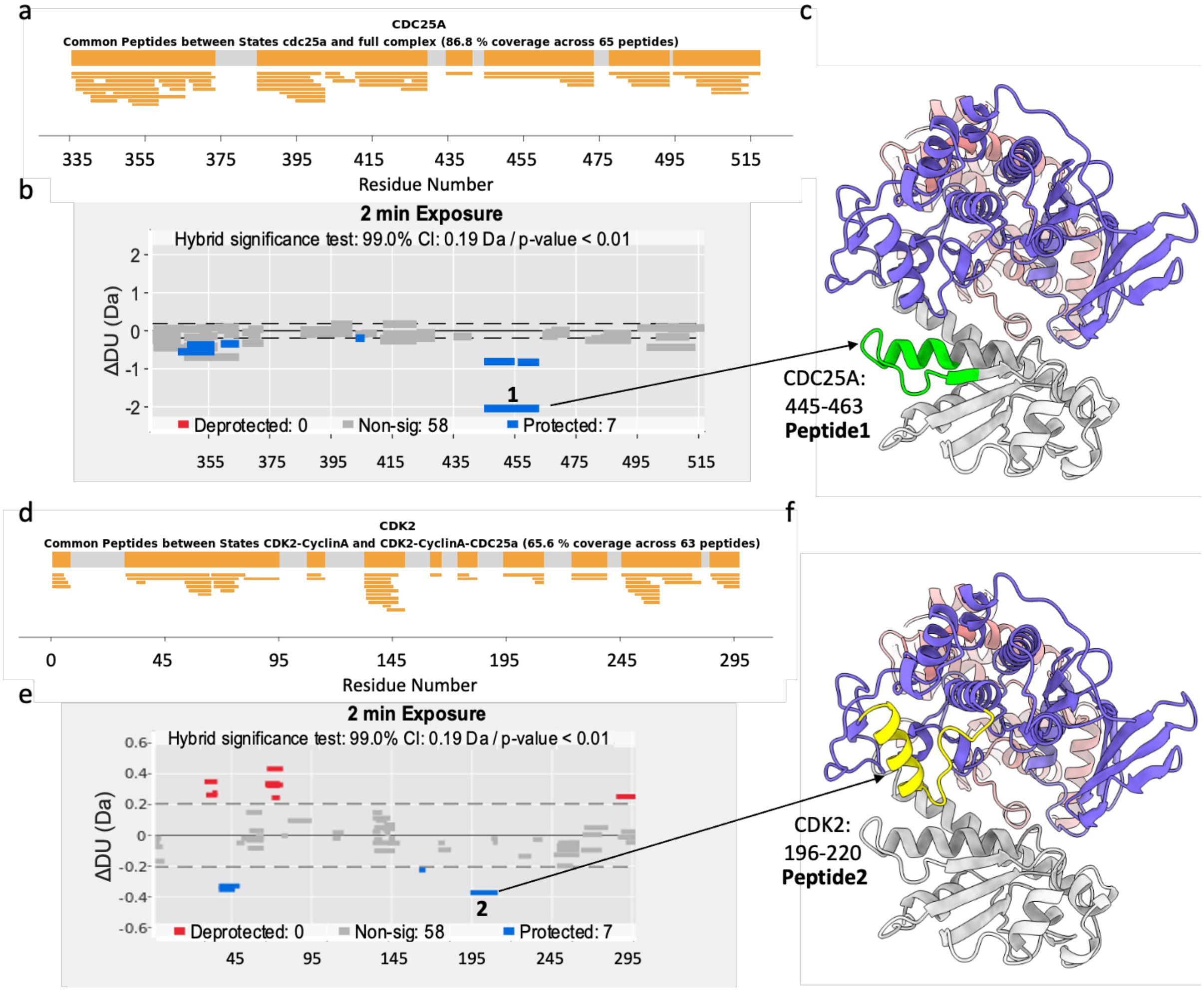

**Supplementary Figure 5: HDX analysis of the CDK2-cyclin A-CDC25A complex**

Peptide coverage plot for (a) CDC25A and (c) CDK2. (b,e) Peptide difference plots for the 2 min deuterium incubation time point for (b) CDC25A and (e) CDK2. The deuterium uptake in the bound state is plotted relative to the unbound state. Peptides highlighted blue show a significant reduction in uptake (protection) in the bound state compared to unbound and red indicates a significant increase in uptake (deprotection). Grey bars indicate no significant difference in uptake between the two states. Locations of (c) Peptide 1 of CDC25A (VRERDRLGNEYPKLHYPEL) highlighted in green and (f) Peptide 2 of CDK2 (MVTRRALFPGDSEIDQLFRIFRTLG) shown in yellow are mapped on the cryo-EM structure. CDK2, cyclin A and CDC25A folds are colored purple, salmon and grey respectively.

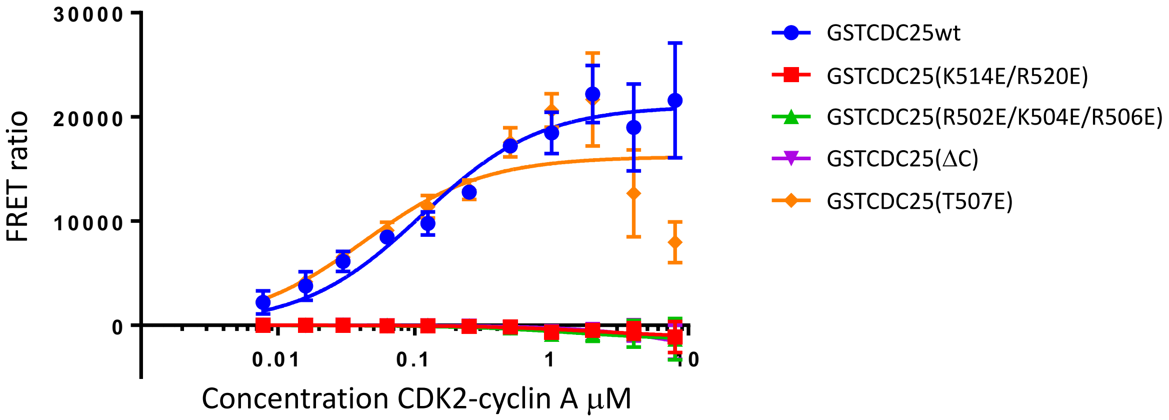

**Supplementary Figure 6: Homogenous time-resolved fluorescence analysis to characterize the CDC25A C-terminal tail interaction with pT160CDK2-cyclin A**

pT160CDK2-cyclin A binding to CDC25A measured by HTRF. The concentration of CDC25A used in these assays was 100 nM. The curves shown are representative binding curves from at least two biological replicates each run in triplicate and carried out on separate days.

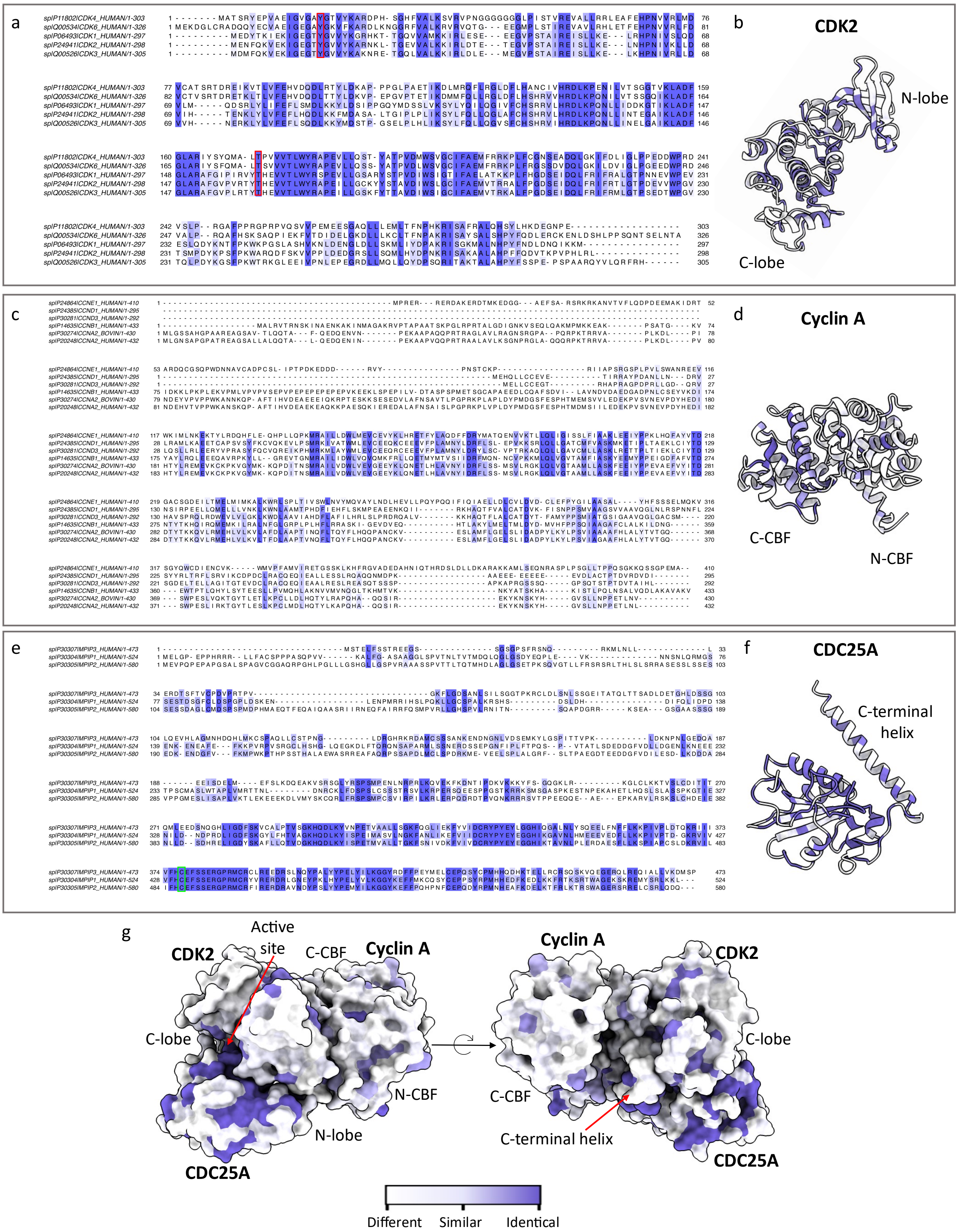

**Supplementary Figure 7: Sequence alignment and conservation analysis of the major cell cycle CDKs, cyclins and CDC25 isoforms.**

Full sequence alignment and conservation analysis of (a) CDK2 vs CDK1, 3, 4 and 6 (c) cyclin A vs cyclin B, E, D1 and D3 and (e) CDC25A vs CDC25B and C. Sequence conservation is indicated in purple (darker shade = identical, lighter shade = similar, white = different). Protein structures (b) CDK2 (d) cyclin A and (f) CDC25A are coloured by sequence conservation. (g) Surface of the CDK2-cyclin A-CDC25A complex coloured by conservation. Sequence alignment was conducted in Clustal Omega followed by conservation analysis and visualisation in Jalview (2.11.2.7). Proteins were coloured by conservation in UCSF ChimeraX.

**Supplementary Table. 1:** Cryo-EM data collection, refinement and validation statistics

| | pY15pT160CDK2-cyclin A-CDC25A (PDB 8QKQ) (EMD-18470) |
|---|---|
| **Data collection and processing** | |
| Magnification | 150,000 |
| Voltage (kV) | 300 |
| Electron exposure ($e^{-}$/Å$^2$) | 50.5 |
| Defocus range (μm) | -2.0 to -0.6 every 0.2 |
| Pixel size (Å) | 0.825 |
| Symmetry imposed | C1 |
| Initial particle images (no.) | 4,981,705 |
| Final particle images (no.) | 670,852 |
| Map resolution (Å) | 2.91 |
| FSC threshold | 0.143 |
| Map resolution range (Å) | 2.4-3.4 |
| **Refinement** | |
| Initial model used (PDB code) | NA |
| Model resolution (Å) | 2.97 |
| FSC threshold | 0.5 |
| Model resolution range (Å) | 2.7 – 3.4 |
| Map sharpening *B* factor (Å$^2$) | -144 |
| Model composition | |
| Non-hydrogen atoms | 6088 |
| Protein residues | 751 |
| *B* factors (Å$^2$) | |
| Protein | 64 |
| R.m.s. deviations | |
| Bond lengths (Å) | 0.004 |
| Bond angles (°) | 0.939 |
| Validation | |
| MolProbity score | 1.32 |
| Clashscore | 5.16 |
| Poor rotamers (%) | 1.1 |
| Fit to map (CCmask, Phenix) | 0.87 |
| Ramachandran plot | |
| Favored (%) | 97.97 |
| Allowed (%) | 2.03 |
| Disallowed (%) | 0.0 |
